# Viscoelastic Notch Signaling Hydrogel Induces Liver Bile Duct Organoid Growth and Morphogenesis

**DOI:** 10.1101/2022.06.09.495476

**Authors:** Muhammad Rizwan, Christopher Ling, Chengyu Guo, Tracy Liu, Jia-Xin Jiang, Christine E Bear, Shinichiro Ogawa, Molly S. Shoichet

**Affiliations:** Department of Chemical Engineering and Applied Chemistry, University of Toronto, Toronto, ON M5S 3E5, Canada; Institute of Biomedical Engineering, University of Toronto, Toronto, ON M5S 3G9, Canada; Terrence Donnelly Centre for Cellular and Biomolecular Research, University of Toronto, Toronto, ON M5S 3E1, Canada; Molecular Medicine Programme, Hospital for Sick Children, Toronto, ON M5G 1X8 Canada; Department of Physiology, University of Toronto, Toronto, ON M5S 1A8, Canada; Department of Biochemistry, University of Toronto, Toronto, Toronto, ON M5G 0A4 Canada; McEwen Stem Cell Institute, University Health Network, Toronto, ON M5G 1L7, Canada; Soham & Shalia Ajmera Family Transplant Centre, Toronto General Research Institute, University Health Network, Toronto, ON M5G 2C4, Canada; Department of Laboratory Medicine and Pathobiology, University of Toronto, Toronto, ON, M5S 1A8, Canada; Department of Chemistry, University of Toronto, Toronto, ON M5S 3H6, Canada

**Author notes:** Corresponding author. M.S. Shoichet.

**Keywords:** Liver organoids, viscoelasticity, mechanotransduction, Notch signaling

## Abstract

Cholangiocyte organoids can be used to model liver biliary disease; however, both a defined matrix in which to emulate cholangiocyte self-assembly and the mechano-transduction pathways involved therein remain elusive. We designed a series of defined viscoelastic hyaluronan hydrogels in which to culture primary cholangiocytes and found that by mimicking the stress relaxation rate of liver tissue, we could induce cholangiocyte organoid growth and significantly increase expression of Yes-associated protein (YAP) target genes. Strikingly, inhibition of matrix metalloproteinases (MMPs) did not significantly affect organoid growth in 3D culture, suggesting that mechanical remodeling of the viscoelastic microenvironment – and not MMP-mediated degradation – is key to cholangiocyte organoid growth. By immobilizing jagged1 to the hyaluronan, stress relaxing hydrogel, self-assembled bile duct structures formed in organoid culture, indicating the synergistic effects of Notch signaling and viscoelasticity. By uncovering critical roles of hydrogel viscoelasticity, YAP signaling and Notch activation, we controlled cholangiocyte organogenesis, thereby paving the way for their use in disease modeling and/or transplantation.

## Introduction

Liver bile duct disorders, collectively known as cholangiopathies, account for up to 70% of pediatric liver transplantations and up to 33% of adult liver transplantations.^1^ Recent breakthroughs in stem cell differentiation and in vitro culture of primary bile duct forming cells (cholangiocytes) have stimulated the development of cholangiocyte organoids for disease modelling, drug testing, and liver tissue engineering.^2,3,4^ Current cholangiocyte organoids depend on reconstituted basement membrane (rBM) surrogates, such as mouse tumor-derived Matrigel, to provide a conducive 3D microenvironment for organoid growth.^2,3,5-9^ Notwithstanding the developments using animal derived rBMs, the cholangiocyte organoids developed to date are rudimentary isolated cysts, which do not undergo morphogenesis into stable bile duct like structures, reflecting limitations of rBMs. Moreover, the rBMs hinder progress in understanding cholangiocyte cell-matrix interactions and mechanisms of cholangiocyte organoid formation due to their ill-defined nature, limited tunability, and batch to batch variability.^10^

Here, we grow primary cholangiocyte organoids in defined hyaluronan hydrogels, and uncover key mechanisms of cholangiocyte mechanosensing in 3D culture, which results in organoid growth. We recapitulate the Notch signaling milieu in a 3D hydrogel and demonstrate that it is required to induce duct-like morphogenesis in cholangiocyte organoids.

Engineered hydrogels in 3D cell culture are often used to model morphogenetic processes in vitro and to elucidate the role of different matrix properties.^11-13^ Matrix stiffness, degradability, cell– matrix adhesion ligands, and ligand density have been shown to impact intestinal organoid formation and budding morphogenesis, neural tube formation, hepatic organoid growth, and kidney epithelial cell lumenogenesis^14,15,16^. More recently, viscoelasticity has been discovered to regulate fundamental cellular processes such as morphology,^17,18^ migration,^19^ and stem cell differentiation^20^. Indeed, many human tissues, including liver, are viscoelastic and display stress relaxation behavior^21^. For example, Chrisnandy et al revealed that a viscoelastic hydrogel facilitates intestinal organoid morphogenesis through dynamic rearrangements of the microenvironment^22^. Indana et al showed that viscoelasticity and adhesion signaling enables human induced pluripotent stem cell proliferation, apicobasal polarization and lumen formation^23^. Surprisingly, the impact of viscoelasticity on cholangiocyte organoid formation, and the molecular events involved in cell-matrix interactions, are largely unknown. Rationally designed matrices capable of providing insights into cholangiocyte morphogenetic processes are key to advancing liver organoid technology.

Liver developmental biology studies and in vitro work revealed that Notch signaling is a key endogenous signaling pathway that controls cholangiocyte differentiation and liver bile duct morphogenesis^24-27^. Portal mesenchyme cells express Jagged1 (Jag1) cell surface ligands, which bind to Notch 2 receptors of neighbouring liver progenitor cells, leading to cholangiocyte differentiation and ultimately bile duct formation^26^. Indeed, co-culture of human pluripotent stem cell (hPSC)-derived liver progenitor cells with the Jag1^+^ mouse stromal cells induced cholangiocyte differentiation, signifying the importance of Notch signaling^28^. The stromal feeder cells were eliminated by incorporating Jag1 into hyaluronan (HA) hydrogels where hPSC-derived liver progenitor cells were differentiated to cholangiocytes in 2D cell culture^29^. Yet, Jag1-induced Notch signaling has not been recapitulated in 3D culture with engineered hydrogels where its role in cholangiocyte organoid morphogenesis can be probed.

HA is a primary ligand of the CD44 receptor and is found ubiquitously in the extracellular microenvironment of various tissues including the liver. HA is distributed in the submucosal space of neonatal and adult extrahepatic bile ducts^30^. Cholangiocytes express CD44 receptors, and hyaluronan is known to increase cholangiocyte proliferation through CD44 binding^31^. We hypothesized that a hyaluronan based 3D hydrogel with tunable composition, degradation, and mechanical properties would enable us to both identify a conducive microenvironment for cholangiocyte organoid growth and elucidate the mechanisms involved in cholangiocyte mechanotransduction.

To test this hypothesis, we designed a new class of stress-relaxing ultra-low content HA-laminin interpenetrating network hydrogels crosslinked using either non-cleavable PEG-tetraoxyamine (PEGOA_4_) or matrix metalloproteinase (MMP)-cleavable peptide bis-oxyamine (MMPOA_2_) (**Fig. 1**). These hydrogels are stable, mechanically tunable, compositionally defined, and suitable for investigating various cell-matrix interactions. We systematically varied hydrogel composition and stress relaxation rates and discovered that HA-laminin hydrogels, with stress relaxation rates matching that of liver tissue, promote primary cholangiocyte organoid growth from encapsulated single cells. This growth is mediated by Yes-associated protein (YAP) signaling and myosin II contractility, but independent of MMP enzymatic remodeling of the matrix. We further modified the hydrogels with Jagged1 using bio-orthogonal norbornene-tetrazine click chemistry, and demonstrated Notch activation in 3D cultured cholangiocytes, which led to unprecedented bile duct-like morphogenesis in 3D culture.

**Figure 1:**
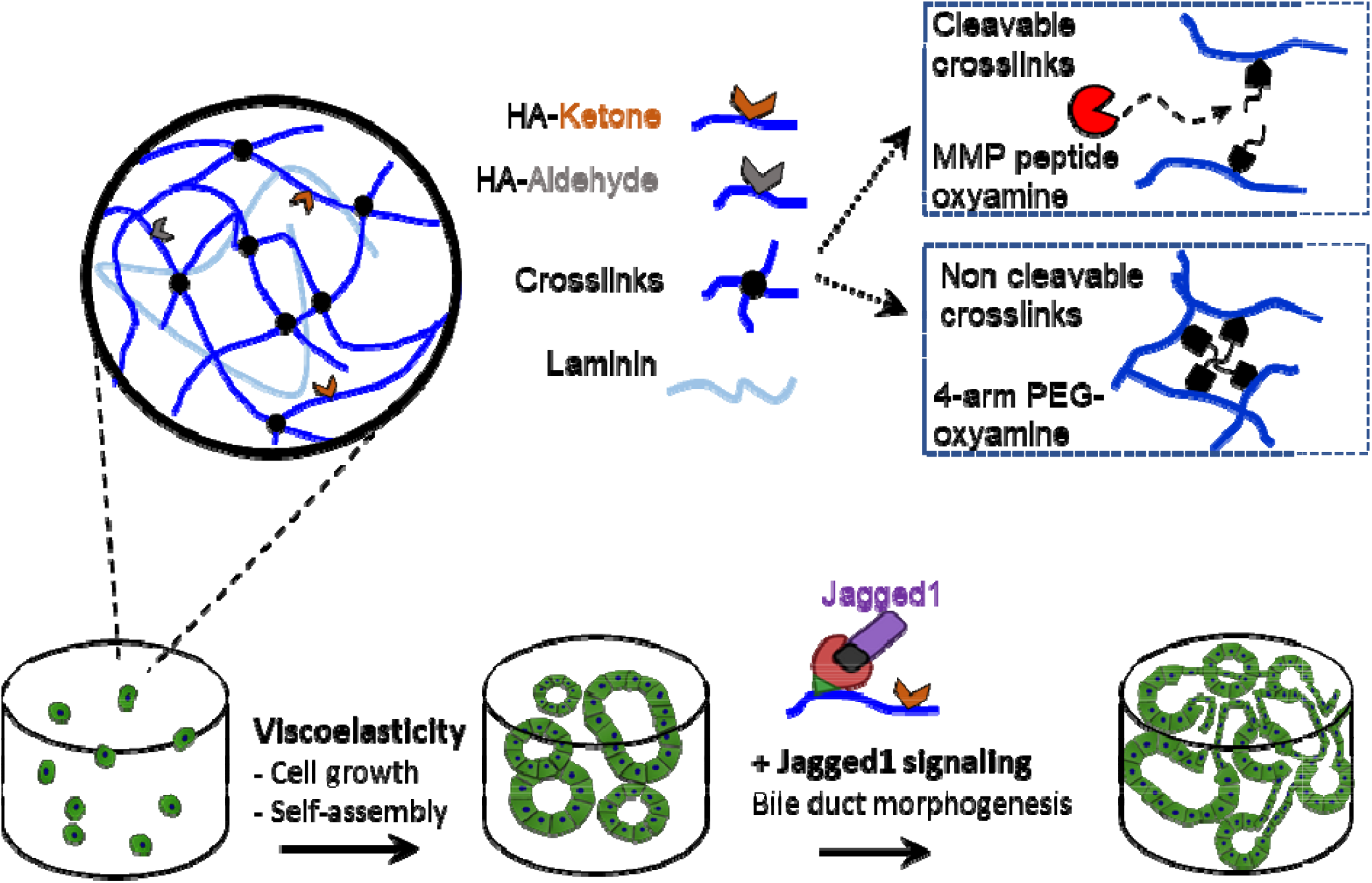
Biochemically and biophysically tunable hyaluronan-based hydrogels (HA) synthesized from fast reacting HA-aldehyde (HAA) and slower reacting HA-ketone (HAK) crosslinked with either non-cleavable PEG tetra-oxyamine (PEGOA_4_) or cleavable matrix metalloproteinase-degradable bis-oxyamine peptide (MMPOA_2_). This hydrogel enables: the growth and polarization of primary cholangiocyte organoids, mechanistic studies of molecular events involved in mechanotransduction, and bile duct morphogenesis through immobilized Jagged1 regulated signalling.

## Results

### Viscoelastic microenvironment stimulates cholangiocyte organoid formation through YAP activation

We investigated the composition of HA oxime (HAO) hydrogels for cholangiocyte organoid growth. HA-ketone and HA-aldehyde were crosslinked using either non-cleavable, oxyamine-terminated 4-arm poly(ethylene glycol) (PEGOA_4_) or cleavable, bis-oxyamine-terminated matrix metalloproteinase-degradable peptide (MMPOA_2_) *OA*-SKAGAGPQGIWGQGAGAKSK(*OA*)S (See Figs S1-4 for characterization of HA-ketone, HA-aldehyde, PEGOA_4_ and MMPOA_2,_ respectively). We found that the incorporation of laminin into HAO hydrogels induced self-assembly of cholangiocytes to organoids (Fig. 2a). As the laminin (Ln) content was increased from 0 to 0.4% (w/v), the cholangiocyte organoid size significantly increased with similar values for 0.3 and 0.4 Ln content hydrogels (Fig. 2b). Thus, we pursued the 0.5% (w/v) HAO-0.3% (w/v) Ln hydrogels where we found organoids with lumens using nuclear and F-actin staining of the cholangiocytes grown therein. In contrast, we only found single cells or cellular clusters in 0.5HAO only gels (i.e., 0.5HAO-0Ln, Fig. 2c). HAO alone did not induce growth of primary mouse cholangiocytes in 3D culture regardless of HA concentration, crosslinking density, and the type of crosslinker used (Fig. S5).

**Figure 2:**
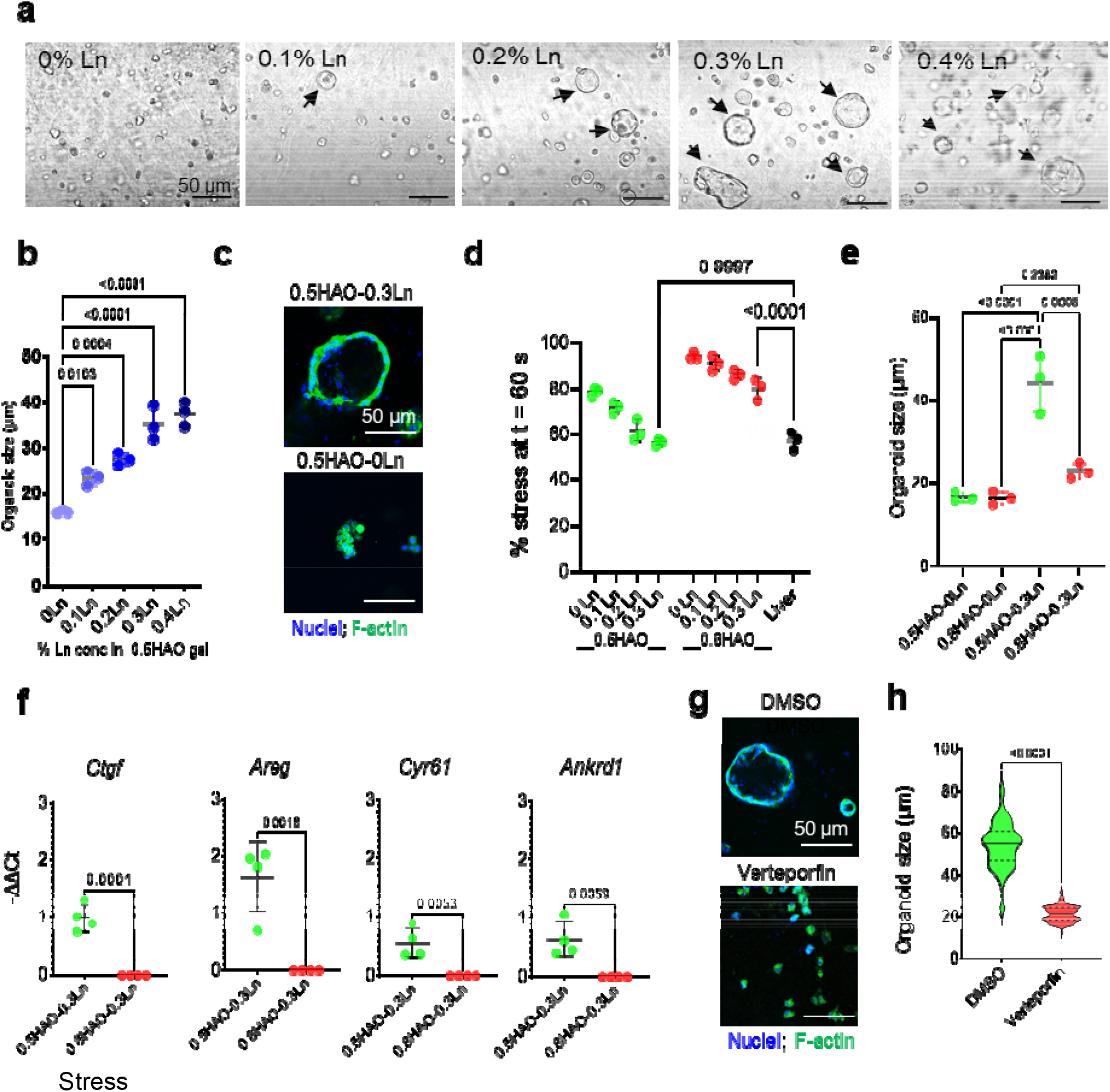
Viscoelastic hyaluronan-oxime laminin hydrogels (HAO-Ln) crosslinked with PEG-oxyamine (PEGOA_4_) provide the microenvironment to stimulate primary mouse cholangiocyte organoid growth, mediated through Yes associated protein (YAP) activation. **(a)** Representative phase contrast images showing the effect of laminin concentration (% (w/v)) on cholangiocyte organoid formation in 0.5% (w/v) HAO hydrogel (arrows highlight organoids). **(b)** Quantification of the effect of laminin concentration on organoid size (*n* = 3 independent experiments, mean ± S.D., one-way ANOVA and Tukey’s post hoc). **(c)** Representative images demonstrating cellular self-assembly to organoids with lumen space in 0.5HAO-0.3Ln hydrogel, which was not observed in 0.5HAO only (with 0Ln) hydrogels. **(d)** Stress relaxation of 0.5% HAO and 0.8% HAO hydrogels with various concentrations of laminin vs. mouse liver tissue: % stress at t = 60 s compared (*n* = 3 hydrogels or liver tissues from 3 separate mouse livers, mean ± S.D., one-way ANOVA and Tukey’s post hoc). **(e)** Effect of HA concentration on the growth of cholangiocyte organoids (*n* = 3 independent experiments, mean ± S.D., one-way ANOVA and Tukey’s post hoc). **(f)** YAP target gene expression in 0.5HAO-0.3Ln (fast relaxing) and 0.8HAO-0.3Ln hydrogels (slow relaxing) (*n* = 4 independent experiments, mean ± S.D., Student’s t-test). **(g**) Representative F-actin-stained images of cholangiocytes cultured in the presence of YAP inhibitor verteporfin or vehicle (DMSO) control. **(h)** Effect of YAP inhibition on the growth of cholangiocyte organoids in fast relaxing 0.5HAO-3Ln hydrogels. Violin plots represent individual organoid sizes (n = 158) from 3 independent experiments. Dotted lines represent quartiles and solid lines are median values. Student’s t-test.

We hypothesized that the HAO-Ln gel induced cholangiocyte growth and self-organization through viscoelastic mechanosensing because laminin is known to form physical crosslinks at higher concentrations, such as those used in our hydrogel. Hydrogels were subjected to a constant strain (15%) with stress relaxation measured as a function of time. Higher laminin content significantly increased stress relaxation rates of both 0.5%HAO and 0.8%HAO hydrogels (Fig S6), with the stress relaxation and percent stress at 60 s of 0.5%HAO-0.3Ln being similar to that of mouse liver tissue (Fig. 2d). To verify that viscoelasticity is the driver of cholangiocyte growth, we cultured cholangiocytes in 0.5HAO-0.3Ln (fast relaxing) and 0.8HAO-0.3Ln (slow relaxing) hydrogels. Strikingly, when the HA content was increased from 0.5% to 0.8%, while keeping the laminin content constant at 0.3%, organoid growth was significantly reduced (Fig. 2e; percent organoids formed, Fig. S7). This highlights the importance of viscoelasticity to organoid growth.

To better understand the role of viscoelasticity, we investigated Yes associated protein (YAP), which is activated during liver bile duct development in vivo and is a key transcriptional element in mechanotransduction. We analyzed YAP activation using well-known YAP target genes (*Ctgf, Cyr61, Areg, Ankrd1*) and observed their significant upregulation in cells cultured in fast relaxing 0.5HAO-0.3Ln compared to slow relaxing 0.8HAO-0.3Ln gels, demonstrating the role of viscoelastic-induced YAP activation in organoid formation (Fig. 2f). Importantly, when YAP was inhibited with verteporfin (5 µM), cholangiocytes remained viable (Fig. S8a) albeit with reduced growth rate (Fig. S8b), but no organoids formed (Fig. 2g). In contrast, cholangiocytes readily formed organoids when cultured in the absence of verteporfin, with vehicle (DMSO) controls, where there were significantly larger organoid diameters compared to cells cultured in verteporfin media, confirming the role of YAP signaling in cholangiocyte organoid formation (Fig. 2h).

### Cholangiocyte organoid growth is dependent on myosin II contractility and FAK-Src signaling, but not on enzymatic remodeling of the matrix

Cellular self-assembly and organoid formation depend on cellular remodeling of the matrix. To understand how the cholangiocytes self-organized to organoids, we investigated a series of mechanisms: focal adhesion kinase (FAK) and Src family kinases (Src), which are downstream of cell-ECM interactions, transduce integrin signals, enhance cell-cell adhesion and promote cell migration; actomyosin contractility; and matrix metalloproteinase (MMP)-mediated digestion / remodeling of the hydrogel microenvironment (Fig 3 for 3D organoid self-assembly; Fig S9 for complementary, 2D cell growth). We investigated whether the FAK-Src complex is involved in cholangiocyte mechanosensing and hence organoid formation. Cholangiocytes encapsulated in 0.5HAO-0.3Ln hydrogels formed significantly smaller spheroids and lacked cellular self-organization when either FAK (with PF-573228) or Src (with dasatinib) was inhibited (Fig. 3a,b). Importantly, neither of the FAK nor the Src inhibitors reduced cell viability (Fig. S9a,b); however, both impacted cell spreading and PF-573228 decreased proliferation relative to vehicle (DMSO) controls (Fig S9b,c). Since the FAK-Src complex is known to induce stress fiber formation and contractility, we treated cholangiocytes with blebbistatin, a myosin II-specific inhibitor, and found that cells were unable to form organoids in 3D culture, demonstrating that actomyosin contractility is critical for cholangiocyte organoid formation from single cells (Fig. 3a,b). Blebbistatin treatment did not reduce cell viability (Fig S9a,b). Collectively, our results indicate that FAK-Src complex, actomyosin contractility, and YAP activation mediate mechanotransduction and enable single cell cholangiocytes to form organoids (Fig. 3c).

**Figure 3:**
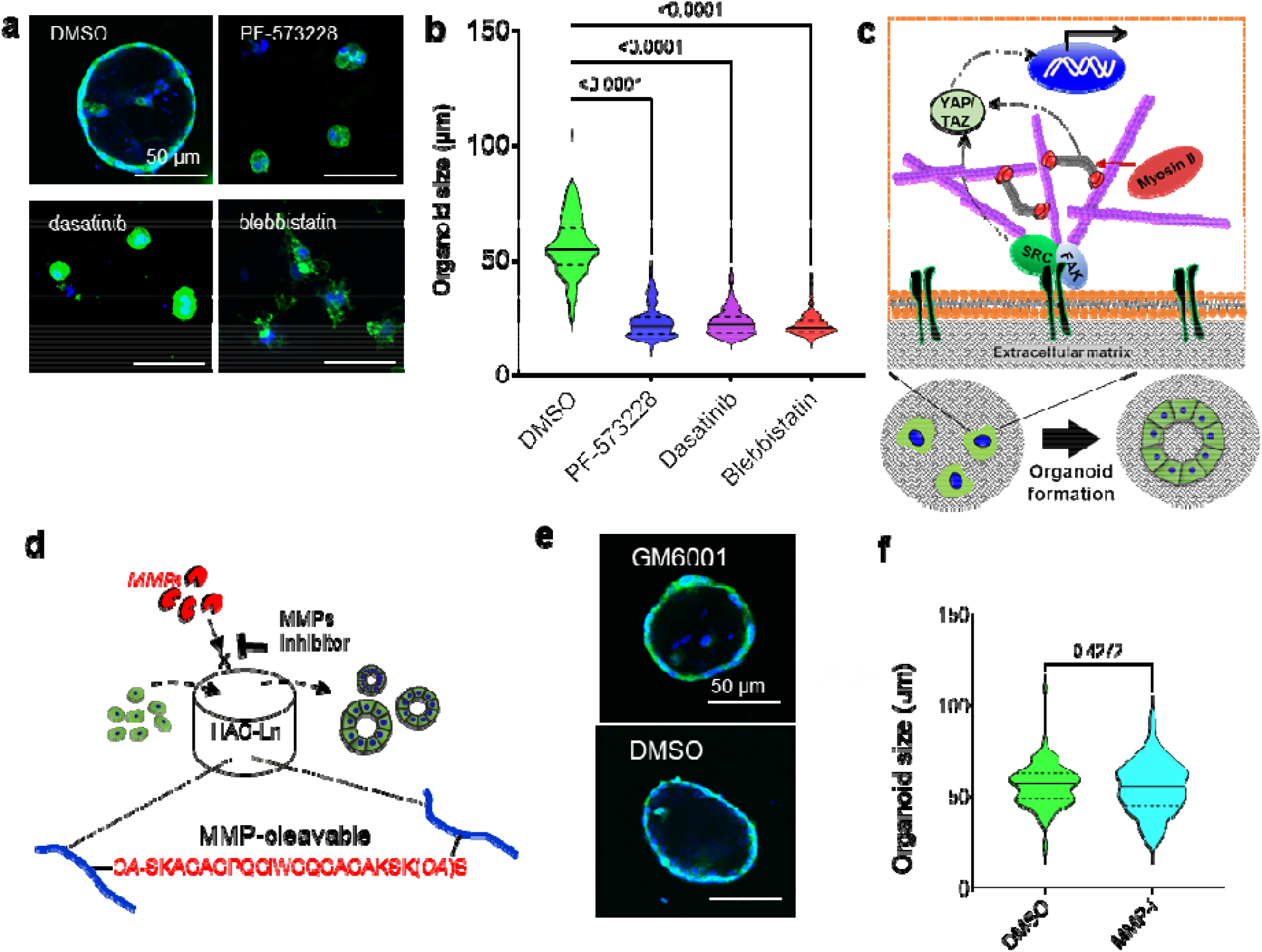
Cholangiocyte organoid growth is dependent on myosin II contractility and FAK-Src signaling, and independent of MMP enzymatic remodeling of the matrix. **(a)** Representative images show the effects of blebbistatin (Myosin II inhibitor), PF-573228 (FAK inhibitor), and dasatinib (Src family kinases inhibitor) on cholangiocyte organoid formation in 0.5HAO-0.3Ln hydrogels at day 5. **(b)** Effect of blebbistatin, PF-573228 and dasatinib on the size of organoids analysed after 5 d of culture. One way ANOVA and Dunnett’s post-hoc. **(c)** Molecular pathways involved in cholangiocyte mechanosensing and transduction leading to organoid growth. **(d)** Schematic of the MMP peptide-cleavable hydrogel used to investigate whether MMP-remodelling of the matrix is involved in cholangiocyte organoid formation. **(e)** Representative images of organoids grown in the presence of GM6001 (pan-MMP inhibitor) or vehicle (DMSO) control. **(f)** Effect of GM6001 on organoid growth in MMP-cleavable hydrogels. Student’s t-test. All violin plots represent individual organoid sizes (n = 152-165 per group) from 3 independent experiments. Dotted lines represent quartiles and solid lines represent median values.

We wanted to understand whether cholangiocytes enzymatically degraded the matrix for growth and self-organization into organoids. To answer this question, we re-formulated our HAO hydrogels using the bisoxyamine MMP-peptide crosslinker (instead of the PEGOA_4_ crosslinker) to form 0.5%HAO-MMP-0.3Ln hydrogels and cultured cholangiocytes therein with or without GM6001 (a pan-MMP inhibitor) (Fig. 3d). Remarkably, single cells were able to form organoids with similar morphology and size even when the MMPs were inhibited (Fig 3e). The size of the organoids in the presence of MMPs was statistically indistinguishable from DMSO controls (Fig. 3f). Importantly, MMP inhibition with GM6001 culture did not affect cell viability (Fig. S9a,b). Cholangiocytes secreted predominantly MMP2 (Fig. S10a) and, when cultured with GM6001, MMP2 secretion was inhibited (Fig. S10b). Thus, cholangiocyte organoid growth is dependent on cellular mechanosensing by the FAK-Src complex, actomyosin contractility, and is independent of enzymatic degradation of the matrix.

### Cholangiocyte organoids express epithelial polarity and biliary markers and are functionally mature

We investigated epithelial polarity and lumen formation in cholangiocyte organoids grown for 7 d in 0.5HAO-0.3Ln hydrogels by staining for nuclei, E-cadherin, β-catenin, and F-actin. F-actin was predominantly present on the outer edges of the cholangiocyte organoids while β-catenin was present on basolateral membranes and E-cadherin was found on lateral cell-cell junctions, confirming cholangiocyte polarization. Immunostaining showed that cholangiocytes expressed mature biliary markers, cytokeratin 19 (CK19) and cytokeratin 7 (CK7), but did not express hepatocyte markers, HNF4α and albumin, demonstrating that cholangiocytes maintain their biliary identity (Fig. 4a).

**Figure 4:**
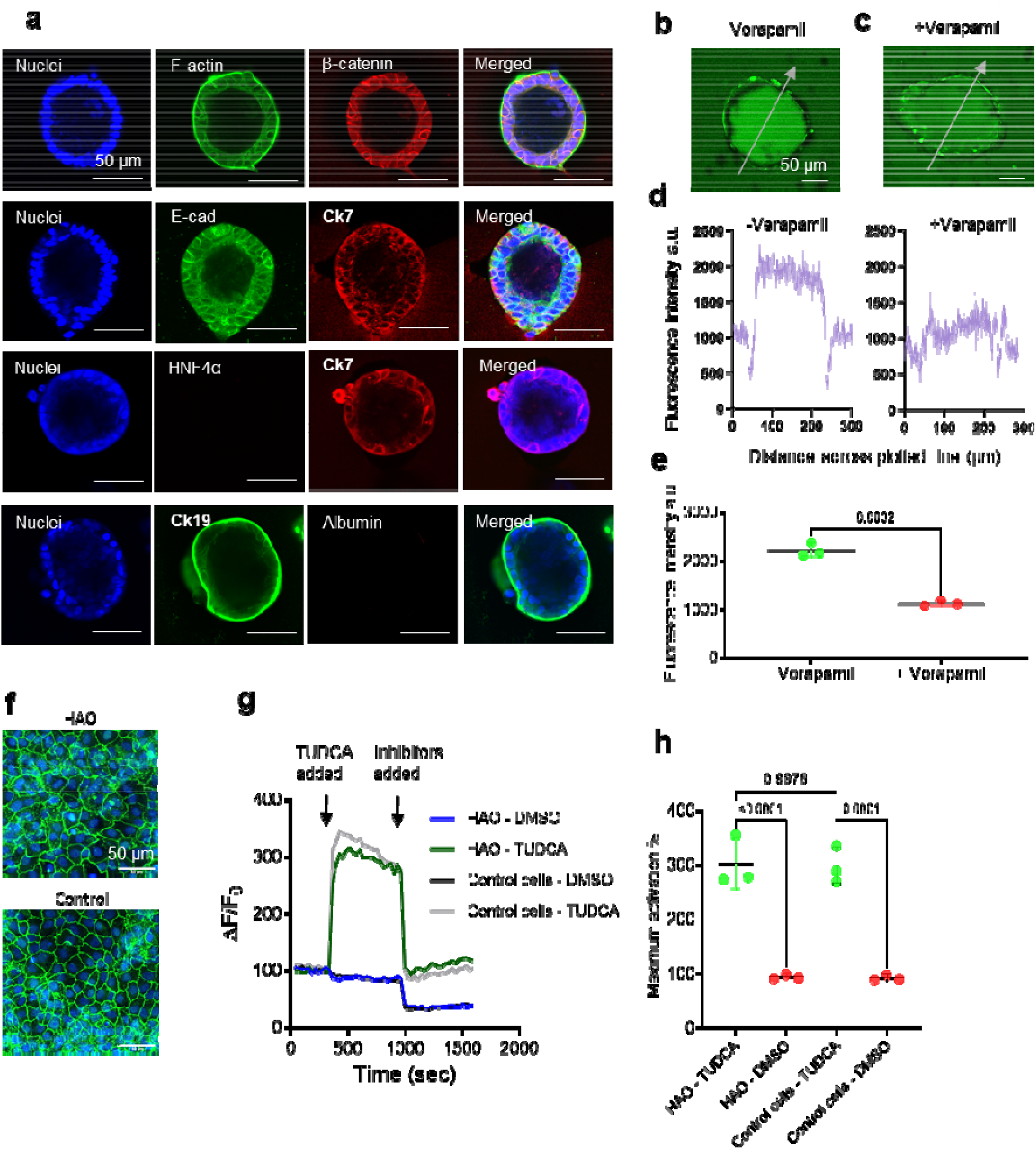
Primary cholangiocyte organoids grown in stress relaxing 0.5HAO-0.3Ln gels demons rate epithelial polarity, express biliary markers and are functionally mature. **(a)** Representative images of the cholangiocyte organoids on day 7 of culture show the localization of F-actin predominantly at the outer edges, β-catenin on basolateral membranes, and E-cadherin at cell-cell junctions. Cells expressed mature ductular markers cytokeratin 19 (CK19) and cytokeratin 7 (CK7) but did not express hepatocytes markers (albumin and HNF4α), confirming their biliary identity. **(b-c)** Rho123 accumulation in the lumenal space of organoids indicating functional MDR1 protein. Verapamil treatment prior to Rho123 incubation inhibited Rho123 accumulation in the lumen of organoids indicating transport is MDR1-specific. **(d)** Fluorescence intensity measurements along the gray lines in b and c. **(e)** Verapamil treatment significantly reduced Rho123 accumulation in the lumen as analyzed by fluorescence intensity measurement of Rho123 (*n* = 3 independent experiments, mean ± S.D., Student’s t-test). **(f)** Cholangiocyte organoids harvested from 0.5HAO-0.3Ln, after hyaluronidase treatment, form monolayers in 2D culture and express ZO1 at cell-cell junctions similar to the positive control cells (grown from passage 1). **(g)** Representative traces of the response of cholangiocyte organoids and positive control cells to the bile acid, TUDCA, and chloride channel inhibitors TMEM16A inhibitor and cystic fibrosis transmembrane conductance regulator (CFTR) inhibitor. Y-axis shows normalized FLIPR® dye signal (F_o_, baseline fluorescence; ΔF, change in fluorescence). Addition of TUDCA caused an increase in chloride conductance leading to membrane depolarization and increase in FLIPR® dye signal. Addition of chloride channel inhibitors reduced chloride conductance, leading to membrane repolarization. **(h)** Maximum % activation relative to baseline shows significantly higher activation in response to TUDCA vs DMSO, and statistically similar activation compared to control cells (*n* = 3 independent experiments, mean ± S.D., one way ANOVA and Dunnett’s post hoc).

In addition to protein marker expression, we evaluated the ability of the organoids to efflux rhodamine 123 (Rho123) into the lumen, which is used to measure multidrug resistance protein 1 (MDR1) activity in normal bile duct cells. Organoids incubated with Rho123 readily transported the dye to the lumenal space, demonstrating the secretory potential of the organoids (Fig. 4b). To confirm that the transport was indeed mediated by MDR1, organoids were incubated with verapamil, an MDR1 inhibitor, prior to Rho123 incubation. Verapamil significantly inhibited lumenal accumulation of Rho123, confirming MDR1-dependent transfer (Fig. 4c,d). We further investigated the functional expression of the apically localized bile acid receptor transmembrane member 16A (TMEM16A) that mediates secretion of bile acids. We first digested the 0.5HAO-0.3Ln hydrogel by incubating it overnight with hyaluronidase (2500 U.mL^-1^), which was tolerated by the cells (Fig. S11). Subsequently, the organoids were plated in a multiwell plate where they formed a monolayer and expressed ZO1 at cell-cell junctions after 1-3 d in culture, similar to cholangiocyte positive control cells grown on tissue culture polystyrene (Fig. 4e). The monolayer was treated with tauroursodeoxycholic acid (TUDCA), a synthetic bile acid. As expected, TUDCA addition induced apical membrane depolarization, measured using the membrane potential dye, FLIPR™. This bile salt-induced depolarization is mediated by apical chloride channels, predominantly TMEM16A. The increase in fluorescence signal caused by membrane depolarization of cholangiocytes was statistically similar to the positive control cells, and significantly higher compared to vehicle (DMSO) control cells (Fig. 4f,g). Importantly, when the cells were subsequently treated with a combination of two inhibitors - a TMEM16A inhibitor and a cystic fibrosis transmembrane conductance regulator (CFTR) inhibitor (CFTRinh-172), the fluorescence signal drastically dropped approaching that of base line fluorescence values, indicating the reversal of TUDCA-induced depolarization (Fig. 4f).

### Notch signaling hydrogel induces bile duct morphogenesis of cholangiocytes organoids

Notch-2 signaling, activated by Jagged1 (Jag1) cell surface ligands, is key to inducing both differentiation of cholangiocytes and development of liver bile ducts; yet, this has only been achieved in 3D culture using OP9 mouse stromal cells. To obviate the use of feeder cells, we developed a new strategy to immobilize full length and bioactive Jag1 protein in 3D 0.5HAO-0.3Ln hydrogels, using a combination of affinity binding and norbornene-tetrazine click chemistry (Fig. 5a). Specifically, protein G-tetrazine (PGtz; mass spectrometry characterization in Fig. S12) was reacted with recombinant Jag1-Fc to obtain the Jag1-Fc-PGtz complex, leveraging the strong Fc affinity binding domains in protein G. Subsequently, the Jag1-Fc-PGtz was reacted with HA-norbornene-ketone (^1^H NMR in Fig. S13), resulting in immobilized Jag1 (Jag1-HAk) that was then mixed with HA-aldehyde, cells, laminin, and crosslinked using PEGOA_4_ to obtain Jag1-0.5HAO-0.3Ln hydrogels. To confirm the successful immobilization and bioactivity of Jag1, we cultured Notch-2 fluorescent reporter cells (N2-CHO cells) and observed significantly greater fluorescence when grown on Jag1-0.5HAO-0.3Ln than on hydrogels without Jag1 (i.e., 0.5HAO-0.3Ln gels) both in 3D (Fig. 5b,c) and 2D (Fig. S14), thus confirming Notch 2 activation. Importantly, simply mixing Jag1 protein fragments (Jag-1 188-204) into the media did not activate Notch 2 (Fig S15).

**Figure 5:**
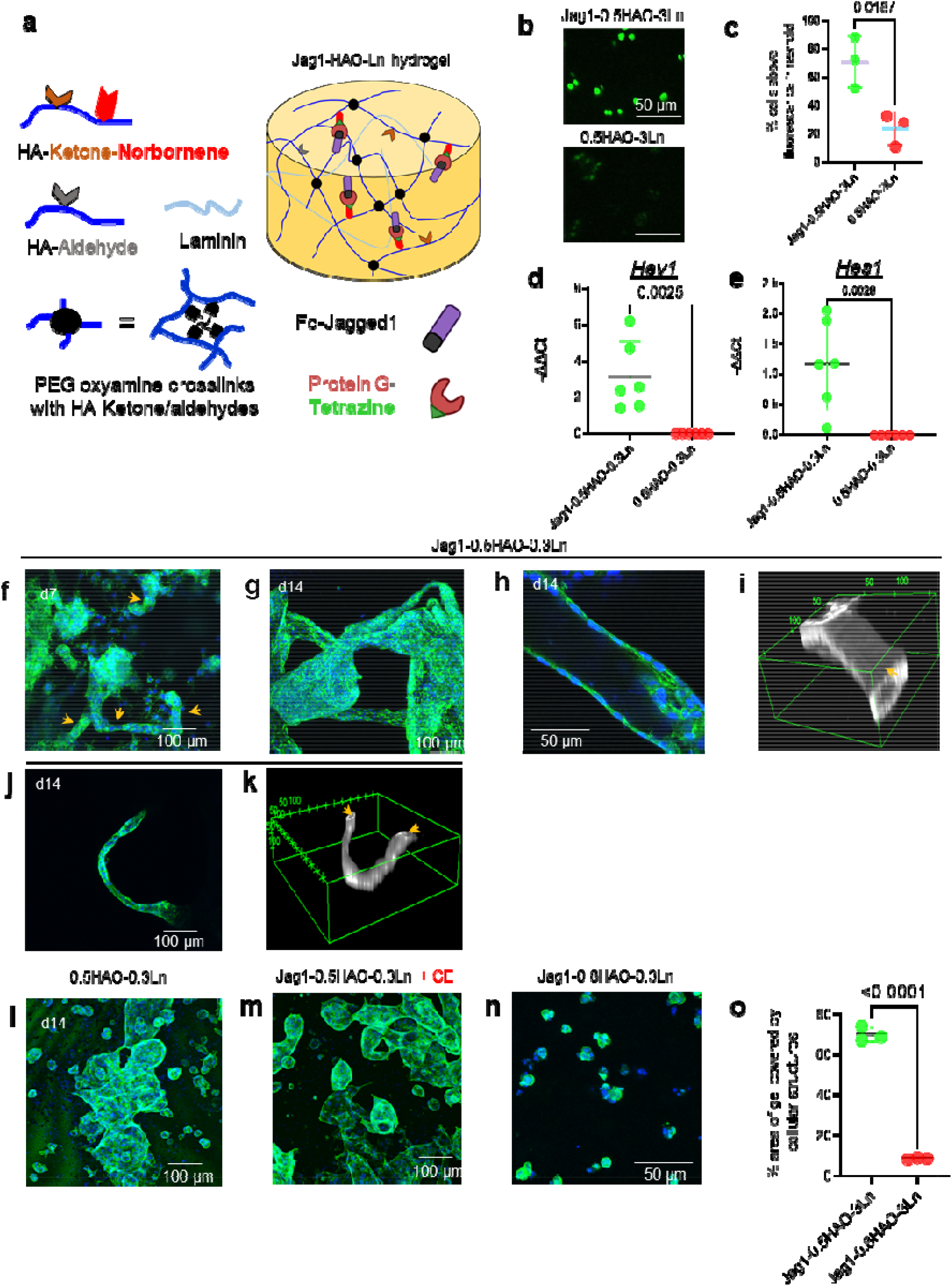
Notch signaling hydrogel induces bile duct morphogenesis of cholangiocytes organoids. **(a)** Formulation of the Jag1-HAO-Ln hydrogel developed to activate Notch signaling in 3D culture of cholangiocytes. (**b,c**) Fluorescent Notch 2 reporter cells (N2-CHO) cultured in Jag1-0.5HAO-3Ln hydrogels show significantly higher fluorescence compared to those cultured in 0.5HAO-3Ln control hydrogels. The fluorescence threshold was set based on the average fluorescence intensity of CHO cells cultured without Jag1. Immobilized Jag1 (on Jag1-0.5HAO-0.3Ln) activates cholangiocyte Notch target gene expression of (**d**) Hey1 and (**e**) Hes1 significantly more than hydrogel only controls (n = 6 independent experiments, mean ± S.D. Student’s t-test for data shown in b-d). Primary cholangiocytes cultured in Jag1-0.5HAO-0.3Ln hydrogels undergo morphogenesis to form duct like structures as early as **(f)** day 7 and **(g)** form larger organoids connected through bile duct-like structures by day 14. **(h,i)** High magnification images of aligned cells forming walls of bile duct-like structures with corresponding 3D rendering of cholangiocyte bile duct, constructed using F-actin-stained cells: the lumen is identified with an arrow (scale is µm for h). **(j,k)** Lower magnification, maximum intensity projection image and corresponding 3D rendering constructed using F-actin-stained cells showing curved bile duct-like structure with lumens observed at both ends (scale is µm for j). **(l)** Cholangiocytes grown in 0.5HAO-0.3Ln hydrogels (in the absence of immobilized Jag1) grew into large, merged organoids without noticeable ductular structures. **(m)** Cholangiocytes cultured in Jag1-0.5HAO-0.3Ln hydrogels in the presence of Notch inhibitor Compound E (CE) do not undergo bile duct morphogenesis. **(n)** Cholangiocytes cultured in Jag1 immobilized slow relaxing 0.8HAO-0.3Ln hydrogels induce neither organoid growth nor morphogenesis. **(o)** Cholangiocytes demonstrate significantly higher growth in Jag1-0.5HAO-3Ln hydrogels than Jag1-0.8HAO-3Ln hydrogels. The cell growth was characterized by analyzing the area covered by cellular structures using maximum intensity projection of Z stacks. (n=3 independent experiments, mean ± S.D. Student’s t-test).

The Jag1-0.5HAO-0.3Ln hydrogel allowed us to study Jag1 induced Notch signaling in a defined 3D environment for ductular morphogenesis of cholangiocyte organoids. We analyzed the Notch activation of encapsulated cholangiocytes cultured in Jag1-0.5HAO-0.3Ln gels for 3 d using quantitative polymerase chain reaction (qPCR). We found that the genetic expression of Notch target genes, Hey1 and Hes1, was significantly higher in cells cultured in Jag1-0.5HAO-0.3Ln gels vs those in hydrogel only 0.5HAO-0.3Ln control gels (Fig. 5d,e).

After the verification of Notch signaling in 3D cultured cholangiocytes, we extended cholangiocyte culture time to 14 d and analyzed morphological changes in cholangiocyte organoids using z-stack confocal microscopy. Fascinatingly, bile duct-like structures appeared in 3D Jag1-0.5HAO-0.3Ln hydrogels connecting cyst like organoids (Fig. 5f). By day 14, cholangiocytes formed larger organoids interconnected through duct-like structures (Fig. 5g) as well as branched duct-like structures (Fig. S16). Higher magnification images showed self-organized cholangiocytes forming the bile duct walls where cell nuclei aligned and elongated laterally to the lumen (Fig. 5h). 3D rendering of the z-stack images using F-actin staining confirmed lumen-containing tubular structure formation (Fig. 5i) and the ability of the cholangiocytes to self-assemble into complex curved bile-duct like structures with lumen (Fig. 5j,k). In contrast, cholangiocytes in 0.5HAO-0.3Ln hydrogels mostly grew to form lumen-containing organoids, which merged resulting in bigger cellular structures (Fig. 5l). To further confirm that the morphological changes are in response to Jag1-induced Notch signaling, we cultured cells in the presence of Notch inhibitor compound E (CE, a small molecule γ-secretase inhibitor). 2D culture studies showed that the CE can inhibit Notch signaling and downregulate Hey1 and Hes1 gene expression without affecting cell proliferation (Fig S17). Single cholangiocytes cultured in Jag1-HAO-Ln gel in the presence of CE (0.1µM) formed round organoids without ductular structures, highlighting the involvement of Notch signaling in ductal morphogenesis, as observed by liver development biology studies (Figure 5m). Finally, we asked whether Jag1 can induce organoid formation or ductal morphogenesis in a slow stress relaxing 3D microenvironment. To study this, we cultured single cholangiocytes in Jag1 immobilized slow relaxing hydrogels (Jag1-0.8HAO-0.3Ln). Cholangiocytes either stayed as single cells or formed small multicellular structures (2-10 nuclei per cluster) after 14 d culture (Fig. 5n). Analysis of the gel area covered by cellular structures using z-stack images showed significantly higher growth of cholangiocyte in Jag1-0.5HAO-0.3Ln hydrogels (fast stress relaxing) vs Jag1-0.8HAO-0.3Ln hydrogels (slow stress relaxing), highlighting that the Notch signaling in a slow relaxing hydrogel is not sufficient to induce organoid growth and morphogenesis (Fig. 5o).

## Discussion

We synthesized hydrogels with defined biochemical and biomechanical environments to provide mechanistic insights into 3D cholangiocyte cellular growth, self-organization, tissue structure and organogenesis. Using defined hyaluronan-laminin based hydrogels, we successfully grew primary cholangiocyte organoids, which has not been previously achieved. Importantly, we identified the role of viscoelasticity and various mechanosensitive pathways, which are critically involved in cholangiocyte organoid formation from single cells, and demonstrated that the synergistic combination of viscoelasticity and Jag1 signaling are key to achieving ductular morphogenesis, thereby recapitulating the Jag1 induced Notch signaling observed in vivo.

Extracellular matrix viscoelasticity has been shown to regulate various cell function and stem cell differentiation;^21^ yet, the role of matrix viscoelasticity has been overlooked in 3D culture of bile duct organoids. The viscoelastic microenvironment can activate YAP signaling,^32^ which is particularly interesting in the context of liver bile duct development where it controls liver cell differentiation^33^ and promotes liver bile duct development in vivo.^34,35^ Our study reveals that the viscoelastic 0.5HAO-0.3Ln hydrogel, with a stress relaxation rate matching that of liver tissue, activates YAP signaling, leading to cholangiocyte organoid growth. Sorrentino et al. found that the Lgr5^+^/EpCAM^+^ liver progenitor organoid formation was also dependent on YAP signaling, but independent of actomyosin contractility.^16^ In contrast, we found that the growth of liver cholangiocyte organoids was significantly impaired when actomyosin contractility was inhibited, highlighting differences in mechanotransduction of liver progenitor cells and liver cholangiocytes. Thus, we hypothesize that actomyosin contractility is required for YAP activation in cholangiocytes. This is supported by previous studies that show significant reduction of nuclear YAP following myosin II inhibition.^36-38^ Moreover, liver progenitor cells require myosin-mediated cell contractility, accompanied by YAP activation, for cholangiocyte differentiation.^39,40^ Interestingly, inactivation of YAP induces defective bile duct morphogenesis due to dysfunction of cholangiocytes.^34^ Our findings complement in vivo studies and confirm the involvement of YAP signaling in bile duct organoid formation.

In cholangiocyte mechanotransduction, we revealed the involvement of the Src-FAK complex that is downstream of cell-ECM interactions. The inhibition of either the Src or FAK pathways abolished organoid formation. The Src-FAK complex mediates integrin signal transduction and connects the extracellular mechanical properties to the actin cytoskeleton. Src-FAK signaling enhances the strength of cell-cell adhesions and promotes cell migration, both of which are key cell functions for epithelial organoid formation.^41,42^ Intriguingly, the activation of focal adhesion-associated kinases, such as Src and FAK, favor stress fiber growth, stability, and contractility, thereby activating YAP.^43^ Taken together, we hypothesize that viscoelasticity sensing is mediated by the Src-FAK complex, which induces actomyosin contractility and YAP activation. YAP signaling ultimately results in cholangiocyte organoid formation. Importantly, our data show that inhibition of any of these pathways results in the loss of cholangiocyte organoid formation. However, it is not clear whether the Src-FAK complex induces organoid formation either directly by modulating cell function or indirectly through YAP activation.^44^ Of note, cholangiocyte organoid growth media often include R-spondin 1 or CHIR99021 that activate Wnt, which, in turn, stimulates the YAP/TAZ pathway. Fascinatingly, we achieved cholangiocyte organoid growth without using a Wnt activator. Thus, YAP activation, induced by the hydrogel microenvironment itself, enabled cholangiocyte organoid growth (in the absence of Wnt signaling), highlighting the importance of cell-ECM interactions in organogenesis.

To gain a greater understanding of how the cholangiocytes remodeled the extracellular microenvironment to form organoids, we investigated viscoelastic hydrogels crosslinked with MMP-degradable peptides (instead of PEG). Interestingly, the cholangiocyte organoids did not require MMP induced enzymatic cleavage of the matrix to form, a trait that was recently discovered in intestinal epithelial organoids as well.^22^ Cholangiocytes formed organoids in non-cleavable but viscoelastic, HA-oxime hydrogels, through structural remodeling of the matrix. The faster stress relaxing 0.5HAO-0.3Ln hydrogel enabled cells to mechanically deform the matrix during cell growth and self-assembly using contractile forces generated by the actin cytoskeleton, which is consistent with the scaffold restructuring reported for other cell types.^22,45-47^ This is further substantiated given that actomyosin contractility is required for force generation^48^ and cholangiocyte organoids did not grow when actomyosin is inhibited.

Our study, for the first time, recapitulates Jag1 induced Notch signaling in organoid culture and demonstrates that cholangiocyte organoids undergo morphogenesis to form duct-like structures. This emulates in vivo studies where Notch signaling is a key driver of bile duct development, thereby validating our models. Remarkably, our data also show that Notch signaling alone does not induce organoid formation if the biomechanical characteristics of the hydrogel are not conducive for cell remodeling. We found that viscoelasticity and Notch signaling work synergistically to provide the appropriate biochemical and biomechanical milieus for organoid morphogenesis. Using engineered HA hydrogels to model the liver environment, we can now manipulate cholangiocyte growth and polarization in a 3D matrix. We understand the mechanosensing pathways involved in the formation of functional cholangiocyte organoids and key factors in organoid and bile duct morphogenesis. We can now use these models to identify new therapeutics and/or targets for liver biliary disease.

## Experimental Section

### Materials

All materials were used as received unless otherwise indicated. Lyophilized sodium hyaluronate (HA; 242 kDa) was purchased from Lifecore Biomedical (Chaska, MN, USA). The following reagents were acquired from Sigma-Aldrich: verteporfin, PF-573228, (-)-blebbistatin, dasatinib, hyaluronidase (type IV-S) from bovine testes, bovine serum albumin, verapamil, Triton X-100, 6-bromohexanoic acid, 10 wt% palladium on carbon, p-toluenesulfonic acid monohydrate, dimethylpropanal, (boc-aminooxy)acetic acid, borane dimethylamine complex, tetrakis(triphenylphosphine)palladium(0), trifluoroacetic acid, N,N-diisopropylethylamine, phosphate buffer saline. The following reagents were used as received: 2,4,6-trinitrobenzene sulfonic acid 5% weight/volume methanol solution (Thermo Fisher Scientific), N-Boc-hydroxylamine (Alfa Aesar), Fmoc-Ser(tBu)-Wang resin (Anaspec), Fmoc-Gly-Wang resin (Anaspec), Fmoc-Lys(Alloc)-OH (Anaspec), 4-(4,6-dimethoxy-1,3,5-triazin-2-yl)-4-methylmorpholinium chloride (TCI), N,N′-diisopropylcarbodiimide (TCI), aminooxy)acetic acid hemihydrochloride (TCI), piperidine (Caledon), methyl cellulose dialysis membrane 12-14 kDa cut-off (Spectrum Laboratories), Jag1 (188-204 (Cedarlane Labs), Hank’s Buffered Salt Solution (Life Technologies), NucleoSpin® RNA purification kit (Macherey-Nagel), SuperScript® VILO™ cDNA Synthesis Kit (Invitrogen), LightCycler® 480 SYBR Green I Master (Roche), laminin-1 (Corning, 354259), rat tail collagen I (Corning), cholangiocyte media and C57BL/6 Mouse Primary Cholangiocytes (Cell biologics, C57-6044C) Nucleospin RNA purification kit (Macherey-Nagel), DNAse (Thermo Scientific), SuperScript III reverse transcriptase (Invitrogen), LightCycler 480 SYBR Green I Master (Roche), goat serum (Gibco), mouse monoclonal anti-CD44 antibody (Abcam, ab6124), all secondary antibodies and Pierce™ protein G coated wells (Thermo Fisher Scientific), DAPI (Roche), Recombinant protein G (Cedarlane Labs Inc). All solvents were purchased from Caledon. Primary antibody sources are given in supporting information Table S2. Fc-Jagged1 was purchased from Sino Biological (Beijing, China).

### Synthesis of HA ketone, HA aldehyde and Star-PEG-tetra(oxyamine) (PEGOA_4_)

Synthesis of HA ketone, HA aldehyde and Star-PEG-tetra(oxyamine) (PEGOA_4_) is described in supporting information.

### Synthesis of oxyamine-modified peptide

*OA*-SKAGAGPQGIQGQGAGKAK(*OA*)S (MMPOA_2_) was prepared using standard Fmoc chemistry using a Liberty Blue peptide synthesizer connected to a Discover microwave (CEM). Fmoc-Ser(tBu)-Wang resin (0.5 mmol) was swollen in DMF for 10 min then F-moc deprotected, and washed using the Liberty Blue peptide synthesizer. A solution containing Fmoc-Lys(Alloc)-OH (0.57 g, 1.25 mmol), HBTU (0.95 g, 2.5 mmol), DIPEA (0.52 mL, 3 mmol) in 5 mL of DMF was added to the resin and left to stand for 10 min. An additional 5 mL of DMF was added and a microwave coupling was performed. The coupling of the Fmoc-Lys(Alloc)-OH on the Ser(tBu)-Wang resin was confirmed by colorimetric TNBS test and the remaining sequence was synthesized. Resin was transferred from the reaction vessel and was washed and then stirred in CH_2_Cl_2_ (8 mL) under nitrogen. Borane dimethylamine complex (20 mg, 0.34 mmol) and tetrakis(triphenylphosphine)palladium(0) (45 mg, 0.04 mmol) were added to the resin suspension and the reaction was stirred overnight protected from light. The solution was drained and the resin was washed with CH_2_Cl_2_ followed by DMF. Resin was treated with 5 mL of Fmoc deprotection solution containing 1 mL piperidine and 1 M 1-hydroxybenzotriazole hydrate (HOBt) for 2 h before being washed with CH_2_Cl_2_. 6-(((tert-butoxycarbonyl)amino)oxy)hexanoic acid (0.30 g, 1.2 mmol) and DIC (193 mg, 1.5 mmol) were combined in 4 mL of CH_2_Cl_2_ for 1 h before adding the solution to the resin. After 24 h, the solution was removed and the resin was washed with CH_2_Cl_2_ and dried. The resin was treated with trifluoroacetic acid (7.2 mL), triisopropylsilane (0.26 mL), phenol (0.26 mL), tryptophan (0.51 g), water (0.26 mL) and aminooxy)acetic acid hemihydrochloride (0.50 mg) for 3.5 h. The peptide was precipitated in cold diethyl ether (50 mL) and washed two additional times. Peptides were purified by column chromatography using C18 silica (SiliCycle) and HPLC using water and acetonitrile containing 0.1% trifluoroacetic acid. Flow rate and gradients were regulated by a 1525 binary HPLC pump (Waters) connected to a xBridge BEH C8 OBD prep column, 5 μm, 19 mm x 150 mm. Fractions containing peptide were identified using a 2489 UV/Vis detector (Waters) and samples were collected using a fraction collector III (Waters). After removing acetonitrile under airflow, fractions were concentrated by lyophilization analyzed by mass spectroscopy. After purification, MMPOA_2_ was obtained as an off-white crystalline solid (326 mg, 30% yield). ESI calculated for C_95_H_157_N_29_O_31_ [M]+: 2201.16; found 2200.16. Purified peptides were stored at -20 °C in glass vials and reconstituted in a 1:1 v/v mixture of 0.1 M sodium phosphate buffer at pH 8 and PBS which were stored in silanized glass vials at 4 °C.

### Synthesis of HA-Norbornene-Ketone (HAnk)

HAnk was prepared in two steps. The first step involved modification of HA with norbornene. 1% w/v sodium hyaluronate (1.247 mmoles, 309 kDa) was dissolved in 2-(N-morpholino)ethanesulfonic acid (MES) buffer (0.1 M, pH 5.5). DMTMM (0.623 mmol) was added to HA solution. After 20 min, 5-Norbornene-2-methylamine (0.623 mmol) was added and stirred for 24 h at room temperature. The solution was dialyzed in 0.1 M NaCl using methyl cellulose dialysis membrane (12-14 kDa molecular weight cut-off) for 24 h followed by distilled water for 24 h, and lyophilized to produce norbornene-modified hyaluronan powder. In the second step, HA norbornene was redissolved in MES buffer, and processed further to modify with ketones as described in supporting information section “Ketone-modified hyaluronan (HAk)”.

### Testing of hydrogel and mouse liver mechanical properties

Hydrogel compression moduli were measured using cylindrical hydrogels with a diameter of 6 mm and thickness of 500 µm with laminin at different concentrations. Hydrogel samples crosslinked using PEGOA_4_ were prepared in a PDMS mold and allowed to crosslink for 2 h at 37 °C and then equilibrated in PBS prior to analysis. Hydrogels were placed between two flat platens connected to a single axis load cell (150 g, ATI Industrial Automation) attached on a Mach-1 micromechanical system (Biomomentum) controlled by a Universal Motion Controller (Newport). An initial force of 0.01 N was used to determine the surface of the sample and the gel height from the distance between the two platens. To remove surface defects a uniaxial unconfined compression was first applied at 10% strain based on the gel height. Samples were tested by applying a further 10% strain. The slope of resultant stress-strain curve for each hydrogel sample was used to calculate the compressive modulus. Experiments were performed with a minimum of 3 independent samples to calculate the mean and standard deviation of compressive modulus. To measure the stress relaxation, hydrogel samples were prepared as described above. The samples were tested by applying a 15% strain and the stress was monitored at 10 Hz over 30 min. The stress values were normalized to each sample.

To measure the stiffness and stress relaxation of the liver tissue, mouse liver tissue slices of 500 µM thickness were prepared using vibratome. Circular disk of 6 mm diameter was trephined from the liver slice to measure the stiffness and stress relaxation as described for hydrogel samples. Samples from 3 mouse livers were used to measure stress relaxation.

### Modification of protein G with tetrazine

5 mg of protein G was dissolved in PBS at room temperature. Methyltetrazine-NHS Ester (1 mol eq to amines) was added to protein G solution, stirred for 15 min at room temperature (RT), and then overnight at 4 °C. Subsequently, the solution was purified with a 7 KDa spin desalting column (ThermoFisher Scientific; Waltham, MA, USA) to obtain tetrazine-functionalized protein G (PG_Tz_), which was characterized by using mass spectroscopy (ESI-MS).

### Primary mouse cholangiocyte expansion and 3D culture in hydrogels

Primary mouse cholangiocytes isolated from extrahepatic liver bile duct tissue of C57BL/6 mouse were purchased from Cell Biologics (C57-6044C). The cells were cultured in epithelial cell medium (M6621) supplemented with Epithelial Cell Medium Supplement Kit (0.5 mL Insulin-Transferrin-Selenium (ITS), 0.5 mL EGF, 5 mL L-glutamin, 5 mL Antibiotic-Antimycotic Solution and 10 mL fetal bovine serum). The cells were passaged with trypsin EDTA when 80-90% confluent. All experiments were performed with cells between passage 3-5. HAk (0.25 – 1.00 wt%) and HAa (0.25 wt%) were mixed together along with cholangiocyte single cells (2 million cells.mL^-1^) and Laminin/entactin (0 - 4.0 mg.mL^-1^) followed by addition of PEGOA_4_ at a 0.25 - 0.75 oxyamine to ketone plus aldehyde ratio to crosslink the HA hydrogel. To prepare enzymatically cleavable HAO hydrogel, MMPOA_2_ was used to crosslink the hydrogel at 0.25 - 0.75 oxyamine to ketone plus aldehyde ratio. The gels were allowed to crosslink for 2 h followed by the addition of cell culture media to the wells. The cells were cultured for 3 to 14 d depending on the experiment as indicated in the results. 50% of the cell culture media was replaced every 2 d.

### 3D cell culture in Notch signaling hydrogel

In a typical experiment, 27.4 µL Fc-Jagged1 (Jag1, 0.5 mg.mL^-1^ in PBS) and 5.2 µL of PGtz (5 mg.mL^-1^ in PBS) were mixed together and incubated at room temperature for 20 min to allow PGtz and Jag1 affinity binding. Subsequently, 100 µL HAnk (25 mg.mL^-1^ in DMEM) solution was added to the PGtz-Jag1 mixture and incubated for 1 h at 37 °C to facilitate norborne-tetrazine reaction. Afterwards, Jag1-PGtz-HAnk was mixed with HAa, cells (2 × 10^6^.mL^-1^) and laminin-1, followed by the addition of PEGOA_4_ crosslinker to initiate gelation. Cells were cultured for up to 14 d in Jag1-0.5HAO-0.3Ln, Jag1-0.8HAO-0.3Ln, and 0.5HAO-0.3Ln hydrogel followed by fixing, permeabilization and staining with Hoechst and Alexa fluor 488 phalloidin overnight as described below. To investigate the effect of Notch inhibition, cells were cultured in Jag1-0.5HAO-0.3Ln hydrogels in the presence of gamma secretase inhibitor compound E, (0.1µM) for 14 d. For the control gels, DMSO was used instead of compound E.

### Gene expression analysis using Quantitative Real-Time PCR (qRT-PCR)

For Hey1 and Hes1 gene expression analysis, cells were encapsulated in Jag1-0.5HAO-0.3Ln and 0.5HAO-0.3Ln gels. After 3 d of incubation, gels were washed for 15 min with PBS and transferred to a centrifuge tube. 350 µL of NucleoSpin RNA extraction buffer along with 4-5 ceramic beads (Qiagen 13113) were added 14into the tube. The centrifuge tubes were placed in a bead beater to homogenize for 1 min, placed on ice for 5 min, then homogenized again for 1 min. Following homogenization, RNA was purified using the NucleoSpin RNA isolation kit as per the manufacturer’s instructions. RNA was reverse transcribed to prepare cDNA using the superscript VILO cDNA kit. Quantitative polymerase chain reaction (qPCR) amplification was performed in an Applied Biosystems 7900HT instrument (40 cycles) using Taqman Assay Kits with 5 ng RNA. The Ct values were normalized to *Gapdh* and cells cultured in 0.5HAO-3Ln hydrogels (-ΔΔCT, -log 2 gene expression). Negative controls (wells without template RNA) were included to ensure accuracy of the data. The primer pairs used are given in Table S1 in the Supporting Information. For YAP target gene expression analysis, cells were cultured for 14 d in 3D 0.5HAO-0.3Ln and 0.8HAO-0.3Ln hydrogels. Subsequently, the RNA was isolated and cDNA was prepared as described above. The qPCR amplification was performed in an Applied Biosystems 7900HT instrument (40 cycles) using LightCycler 480 SYBR green with 5 ng RNA. The Ct values were normalized to *Gapdh* and cells cultured in 0.8HAO-0.3Ln hydrogels (-ΔΔCT, -log 2 gene expression). Negative controls (wells without template RNA) were included to ensure accuracy of the data. The primer pairs used are given in Table S1 in the Supporting Information.

### Immunofluorescence staining of cholangiocytes

For immunostaining following culture, the cells/organoids were fixed with 4% paraformaldehyde for 60 min. After three washes in PBS (30 min each), cells were permeabilized with a mixture of 100 × 10^−3^ M glycine and 0.5% triton X-100 for 1 h, washed with PBS, and blocked with 10% goat serum and 1% bovine serum albumin (BSA) overnight. Following blocking, organoids were incubated overnight with primary antibodies, washed with PBS (3 times, 1 h each), and incubated with secondary antibodies overnight. List of primary and secondary antibodies and their dilutions are provided in Table S2. Subsequently, the cells were washed again with PBS (3 times, 1 h each) and stained with 20 × 10^−3^ M Hoechst 33342 dye for nuclei staining and Alexa Flour 488 phalloidin for F-actin staining. For protein expression analysis, following staining the organoids were retrieved from hydrogels by incubating overnight in hyaluronidase solution (5000 U.mL^-1^ in PBS) for improved imaging resolution. For the analysis of organoids morphogenesis in Jag1 immobilized hydrogels, the imaging was performed in 3D hydrogels. Organoids were imaged with an Olympus FV1000 confocal microscope.

### Characterization of organoid formation and growth in 3D hydrogel culture

Brightfield Z stack images of the cells/organoids were captured at day 0 after cell encapsulation (single cell stage) and day 3 (organoid formation) using Olympus FV1000 confocal microscope at different depths in the hydrogels (10 µm step size). Ferret’s diameter was measured using the “analyze particle” function in ImageJ to quantify the size of organoids formed from single cells in hydrogels with different concentrations of laminin or HA. The particles below 15 µm diameter (threshold set using analysis of single cell size after encapsulation) were excluded from analysis. Images from 3 independent replicates were used to analyze organoid size. Between 150 – 200 organoids were analyzed (>50 per replicate).

To analyze the cell morphology and organization in organoids, cells were stained with Hoechst 33342 and Alexa Fluor™ 488 phalloidin overnight as described above, and imaged using confocal microscopy. To analyze organoid morphogenesis in response to Jag1, the cellular structures were imaged using confocal microscopy after 14 d culture. Z stacks images were collected through the hydrogel thickness with 4 - 20 µm step size at 5 different locations in the gels. ImageJ was used to prepare 3D rendering of the tubular structures using Alexa Fluor™ 488 phalloidin channel, maximum intensity projection images, z stack animations, and to quantify the area covered by the cellular structures as a proxy for cell growth. Z stack animations (4 frames per second) represent 20-25 images captured with 10 µm step size through the thickness of the gel to demonstrated 3D structure of organoids and ductular structures. For % area coverage analysis, maximum intensity projection images from 3 independent replicates were analyzed in each group.

### Effect of various pathway inhibitors on organoid formation

To investigate the effect of various signaling pathway inhibitors on cholangiocyte organoid growth, primary mouse cholangiocytes were encapsulated in 0.5%HAO-0.3Ln hydrogel crosslinked with PEGOA_4_ and cultured for up to 5 d in the presence of following inhibitors: Verteporfin, Dasatinib, Blebbistatin, and PF-573228 (all at 10 µM concentration, except verteporfin at 5 µM). DMSO was used as the vehicle control. Afterwards, cells were fixed, permeabilized and stained with Hoechst 33342 and Alexa Fluor 488 phalloidin overnight as described above. Images from 3 independent replicates were used to analyze organoids size. Between 150 – 200 organoids were analyzed (>50 per replicate). To investigate the role of MMP induced microenvironment remodeling, cells were cultured with pan MMP inhibitor GM6001 at 10 µM concentration or DMSO for 5 d in 0.5%HAO-0.3Ln hydrogel crosslinked with MMPOA_2_ peptide. Subsequently, the organoids were stained and analyzed as described above.

### Characterization of Notch activation using reporter cells

Notch2 expressing polyclonal CHO-K1 fluorescent reporter cells (N2-CHO cells) were kindly donated by the Elowitz lab (California Institute of Technology).^49^ The cells were cultured in αMEM media supplemented with 10% FBS, penn/strep, 2 mM L-glutamine, and passaged when 80-90% confluent. To analyze the Notch signaling activation, N2-CHO cells were seeded on Jag1-0.5HAO-0.3Ln gels at 1.5 × 10^4^ cells/cm^2^, cultured for up to 3 d and imaged using the confocal microscope. 0.5HAO-0.3Ln hydrogels without Jag1 ligands were used as control.

### Functional testing of cholangiocytes using Rhodamine 123 (Rho123) transport and chloride conductance assay

Cholangiocyte organoids were grown in 0.5HAO-0.3Ln hydrogels for 14 d. Hydrogels were washed 3 times (5 min each) with phenol red–free Hank’s balanced salt solution (HBSS), and then incubated with HBSS containing 100 μM rhodamine 123 for 30 min. Subsequently, the gels were washed again with HBSS (3 times, 5 min each) and the rhodamine accumulation was visualized using a confocal microscope. To inhibit MDR1 transporter, the organoids-containing hydrogels were incubated with 20 μM R-(+)-verapamil (Sigma-Aldrich) for 30 min before the rhodamine 123 assay as described above. ImageJ was used to measure fluorescence intensity in the lumen of organoids with or without verapamil treatment.

To assess CFTR or TMEM16A mediated change in membrane depolarization, the apical chloride conductance assay (ACC) was used as previously described^50,51^. In brief, mouse cholangiocyte cysts were seeded for 48 h and the cultures were incubated in a chloride-bicarbonate buffer without sodium (NMDG 150 mM, Gluconic acid lactone 150 mM, Potassium Gluconate 3 mM, Hepes 10 mM, 0.5mg/mL FLIPR™ dye, pH 7.4, osmolarity 300 mOsm) for 30 min. Chloride channel conductance recordings were measured at 30 s intervals on the Synergy Neo2 Multi-Mode Assay Microplate Reader (BioTek). Following 5 min of baseline recording, cultures were treated with 2.5 mM TUDCA and monitored for 10 min.^52^ Lastly, channel activities were terminated by the addition of CFTRinh172 (10 µM) and inhibitor of TMEM16A (CCAHinh, 10 mM) for 10 min.

### MMP expression analysis

Mouse cholangiocytes were cultured in 6 well plate at seeding density of 1 × 10^5^ cells per well. After 2 d, whole media was removed and fresh media with or without GM6001 inhibitor (10 µM) was added to the cells. After 24 h, the media was collected and frozen at -20 °C. The MMP expression and inhibition was analyzed using zymography as previously described.^53^ Briefly, 6 µL media was mixed with 2 µL loading dye without β-mercaptoethanol. Samples were loaded on Novex 10% Zymogram Plus gelatin protein gels and the gels were run and washed as previously described. Gels were then stained with 0.4% Coomassie Blue for 2 h, followed by destaining (10:30:60 acetic acid: methanol: water, v/v/v) for 30 min, and imaged with an E-gel Imager (Invitrogen). Image J was used to quantify gelatin degradation.

### Statistical analysis

All statistical analyses were performed using GraphPad Prism version 8. Differences between two groups were analyzed using student’s t-test. Differences between more than two groups were analyzed using one-way analysis of variance (ANOVA) and Tukey’s post hoc test. A P-value of <0.05 was deemed statistically significant. Experiments were repeated using cells from a differenty passage number.

## Acknowledgments

We are grateful to the Canada First Research Excellence Fund to Medicine by Design at the University of Toronto for funding.

## Author contributions

M.R. and M.S.S. conceived the project, designed the experiments, interpreted the results, and wrote the manuscript. M.S.S. revised the manuscript. M.R. performed most of the experiments. C.L. performed Notch activation experiments. C.G. and T.L. assisted in characterizing the effect of hydrogel composition on organoid formation and stress relaxation experiments, respectively. J.J. performed chloride channel experiments. S.O. and C.E.B. provided experimental guidance.

