## supporting info for "Viscoelastic Notch Signaling Hydrogel Induces Liver Bile Duct Organoid Growth and Morphogenesis"

**Supporting Information**

**Methods**

**Ketone-modified hyaluronan (HAk)**

HAk was synthesized as previously reported^1^. Briefly, 1% w/v sodium hyaluronate (1 g, 242 kDa) was dissolved in 2-(N-morpholino)ethanesulfonic acid (MES) buffer (0.1 M, pH 6.6). DMTMM (4.36 mmol) was added to HA solution. After 30 min, 3-(2-methyl-1,3-dioxolan-2-yl)propan-1-amine (2.5 mmol) was added and stirred for 48 h. The solution was dialyzed in 0.1 M NaCl using methyl cellulose dialysis membrane (12-14 kDa molecular weight cut-off) for 48 h followed by distilled water for 24 h. Subsequently, the ketal-substituted hyaluronan was dialyzed against 0.2 M HCl for 2 h. The solution was changed to pH 4.0 with sodium bicarbonate solution (0.1 M) overnight and then further dialyzed against distilled water for 48 h. The dialysate was sterile filtered and lyophilized. The degree of substitution was analyzed by ^1^H NMR in deuterium oxide to be 43%.

**Aldehyde-substituted hyaluronan (HAa)**

HAa was synthesized as previously reported^1^. Briefly, 1% w/v HAA was prepared by dissolving sodium hyaluronate (1.00 g, 242 kDa) in MES buffer (0.1 M, pH 5.5), and stirred with DMTMM (4.36 mmol). After 15 min, aminoacetaldehyde dimethyl acetal (1.0 mmol) was added dropwise and stirred at 60 °C for 24 h. The solution was dialyzed in 0.1 M NaCl using 12-14 kDa molecular weight cut-off dialysis membrane for 48 h followed by distilled water for 24 h. Subsequently, the acetal-substituted hyaluronan was dialyzed against 0.2 M HCl for 48 h. The pH of the solution was changed to 4.0 using sodium bicarbonate solution (0.1 M) overnight and then dialyzed against distilled water for 48 h. The dialysate was sterile filtered and lyophilized. The degree of substitution was analyzed by ^1^H NMR in deuterium oxide to be 41%.

**Star-PEG-tetra(oxyamine) (PEGOA_4_)**

PEGOA_4_ was prepared by dissolving (Boc-aminooxy)acetic acid (0.43 g, 2.3 mmol) and *N,N'*-diisopropylcarbodiimide (DIC) (0.57 mL, 4.6 mmol) in 25 mL of dichloromethane (DCM) at 0 °C under nitrogen. After 1 h PEG-tetramine (1.0 g, 5250 Da) was added followed by *N,N*-diisopropylethylamine (1.18 mL, 9.14 mmol), and the reaction was stirred for 48 h at room temperature. DCM was removed in vacuo and the crude was stirred with distilled water and filtered through a 0.22 μm filter to remove the *N*,*N*'-diisopropylurea by-product. The filtrate was then dialyzed using methyl cellulose dialysis membrane (1,000 Da molecular weight cut-off) in sodium chloride (0.1 M) for 24 h followed by distilled water for 48 h before being lyophilized. Boc-protected Star-PEG-tetra(oxyamine) was obtained as a white solid and characterized for substitution by ^1^H NMR before deprotecting by dialysis in hydrochloride acid (0.2 M) for 48 h followed by distilled water for 48 h. The dialysate was lyophilized to give star-PEG-tetra(oxyamine) (PEGOA_4_) as a white crystalline solid (0.95 g, 46.1% yield). ^1^H NMR was performed to determine the completion of deprotection using CDCl_3_.

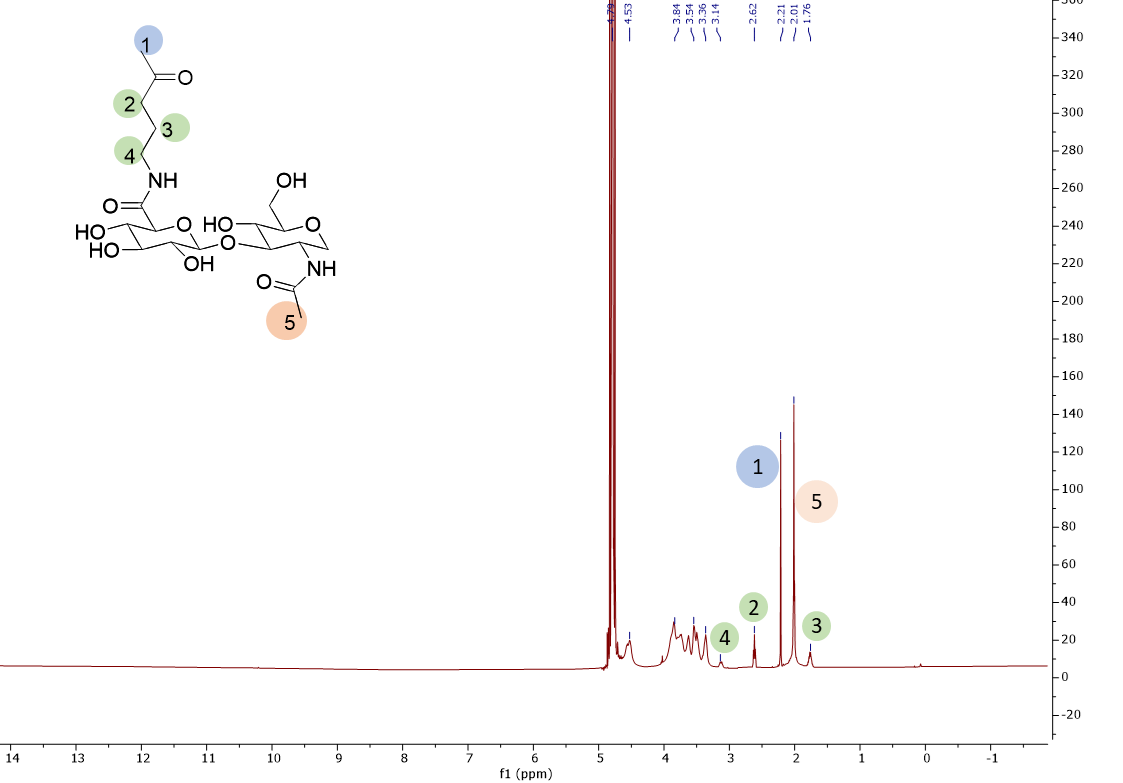

Fig. S1: ^1^H NMR spectrum of HA-ketone in D_2_O.

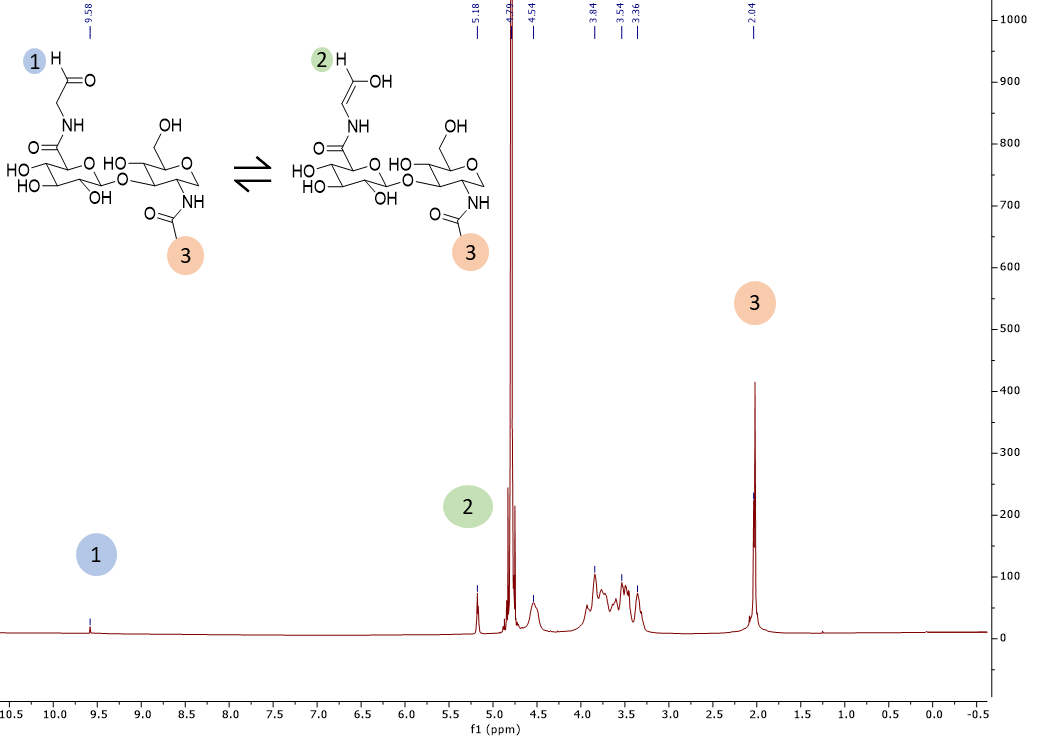

Fig. S2: ^1^H NMR spectrum of HA-aldehyde in D_2_O.

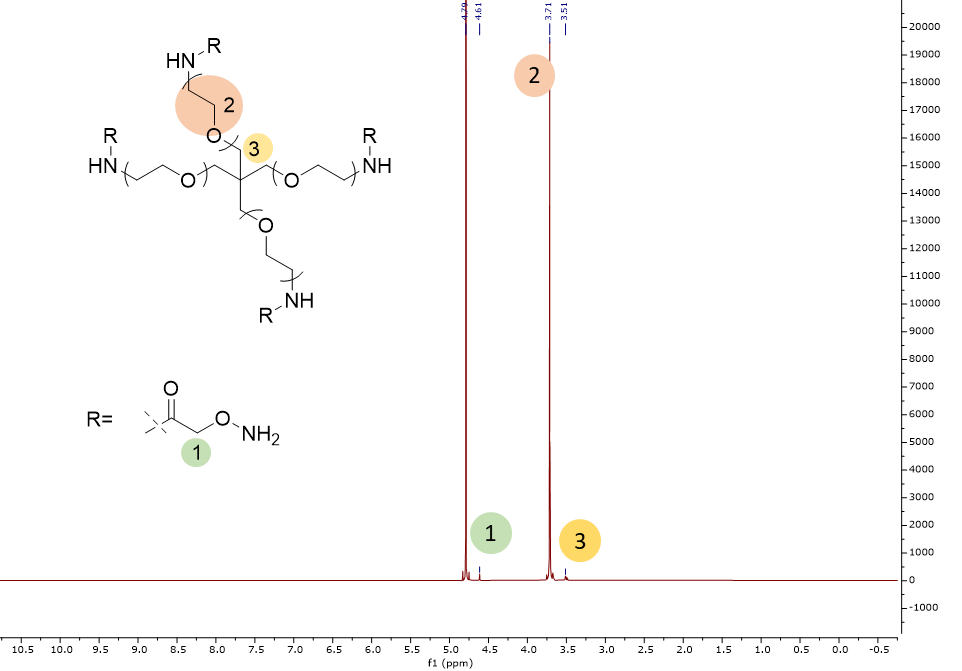

Fig. S3: ^1^H NMR spectrum of 4-arm PEG oxyamine (PEGOA_4_) in D_2_O.

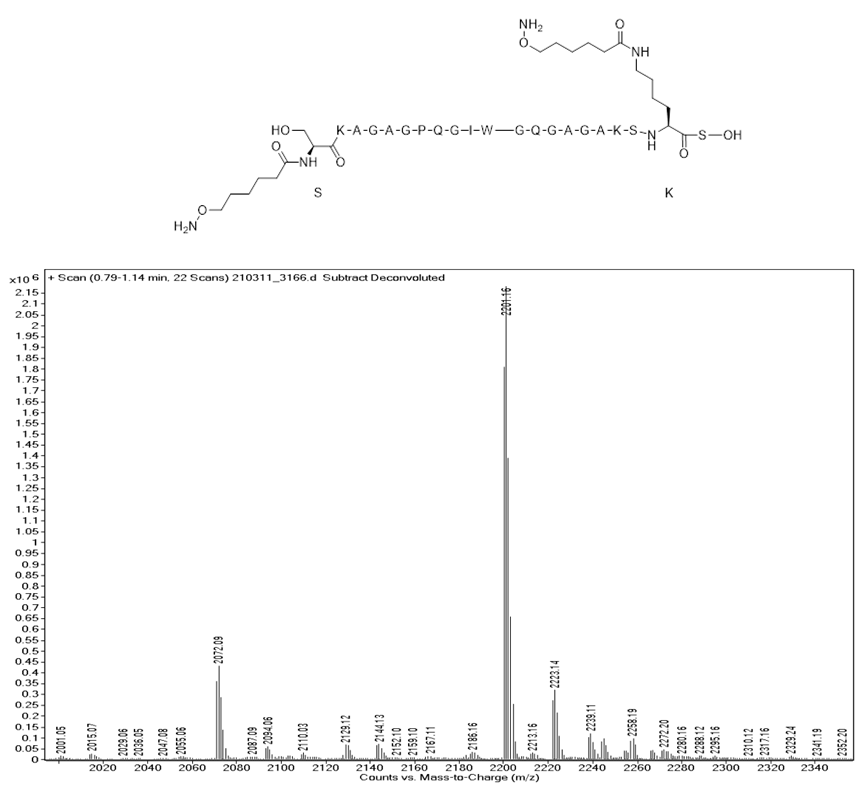

Fig. S4: Mass spec of MMP-cleavable peptide bis-oxyamine (MMPOA_2_): *OA*-SKAGAGPQGIWGQGAGAKSK(*OA*)S. Expected mass for C_95_H_157_N_29_O_31_ [M]+: 2201.47 g/mol; Observed: 2201.16 g/mol.

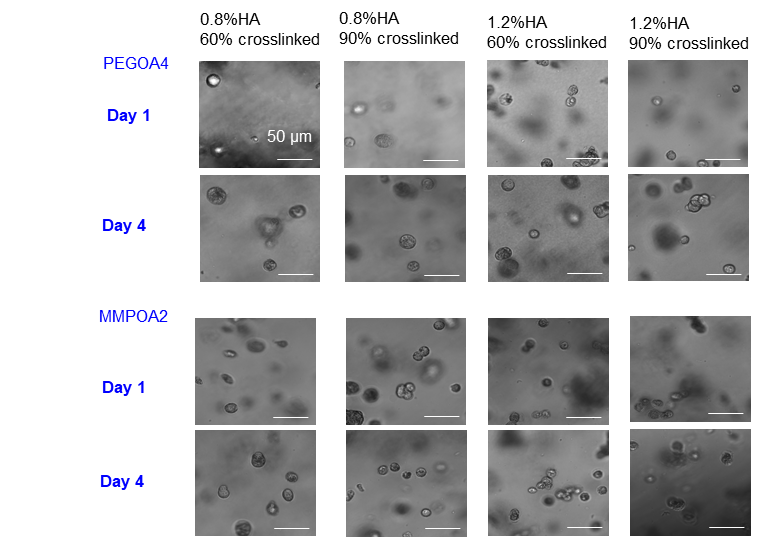

Fig. S5 Primary mouse cholangiocytes neither spread nor proliferated in HA oxime (HAO) hydrogels crosslinked with either PEGOA_4_ or MMPOA_2_ peptides.

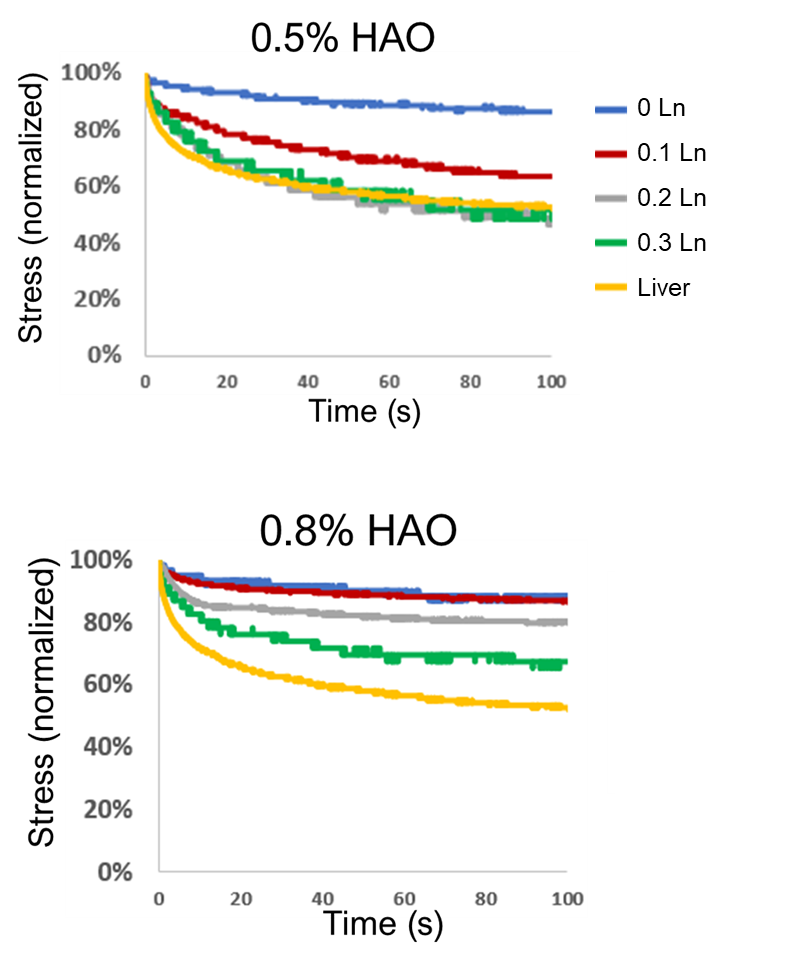

Fig. S6: Representative stress relaxation curves of 0.5HAO and 0.8HAO hydrogels with various amounts of laminin. Samples were subjected to a constant 15% strain and the stress was recorded at various time points, which was then normalized to the stress value at t=0 s (the onset of stress relaxation) for each sample. Mouse liver tissue had a similar stress relaxation profile to that of 0.5HAO-0.3Ln.

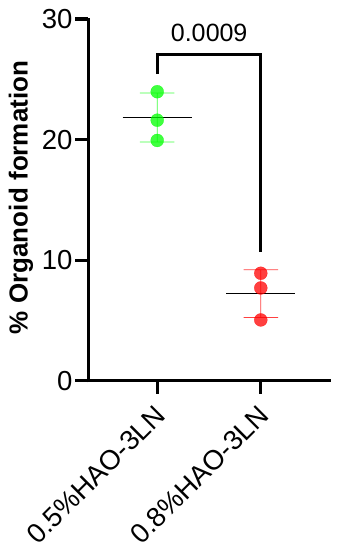

Fig S7. Effect of HA content on the percent of organoids formed measured at day 3 defined as the % cholangiocyte single cells which form organoids. (n=3 independent experiments, mean ± S.D. Student’s t-test.)

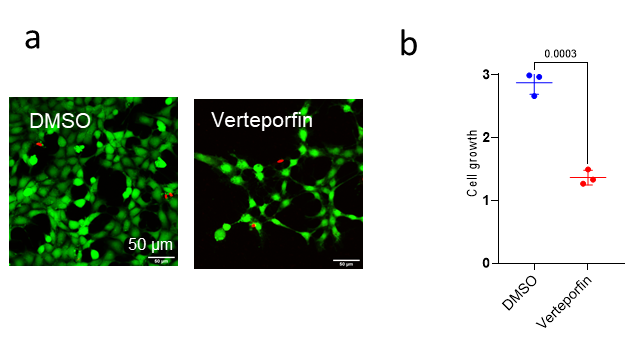

Fig S8: (a) Representative images of primary mouse cholangiocytes stained with Calcein AM (green, live cells) and Ethidium bromide (red, dead cells) at day 3 with or without YAP inhibitor, verteporfin. (b) The PrestoBlue assay was used to quantify the number of cells cultured in the presence or absence of verteporfin. Data shows cells growth at day 3 normalized to day 1. n=3 independent experiments, mean ± S.D. Student’s t-test.

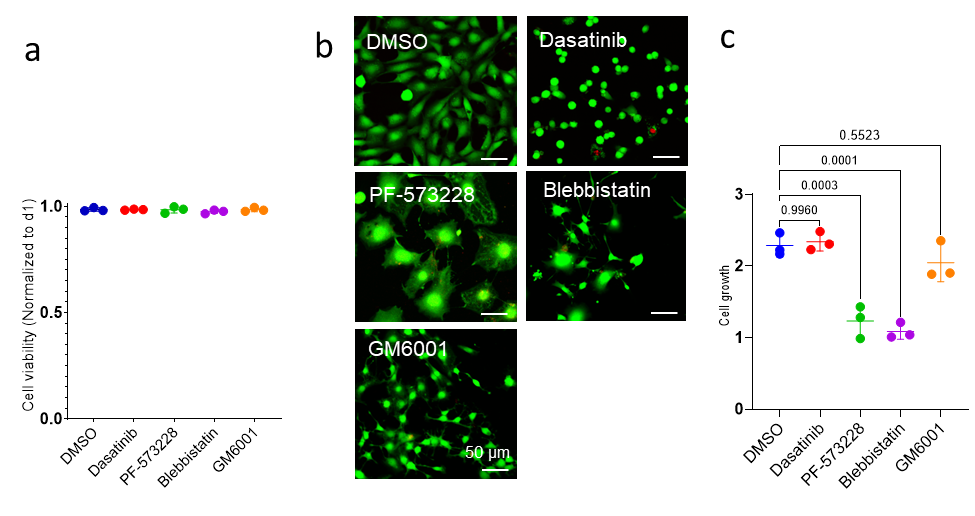

Fig S9: (a) Normalized primary mouse cholangiocytes cell viability at day 3 in the presence of different inhibitors: DMSO is the vehicle control; dasatinib inhibits Src kinases; PF-573228 inhibits FAK; blebbistatin inhibits actomyosin; GM6001 inhibits MMPs. Cell viability was measured by counting the cells stained with Calcein AM (green, live cells) and Ethidium bromide (red, dead cells). The data was normalized to day 1. (b) Representative images of primary mouse cholangiocytes stained with Calcein AM and Ethidium bromide day 3 in the presence of different inhibitors. (c) The PrestoBlue assay was used to quantify cell growth in different conditions. Data shows cell growth at day 3 normalized to day 1. n=3 independent experiments, mean ± S.D. One way ANOVA followed by Dunnett’s post hoc.

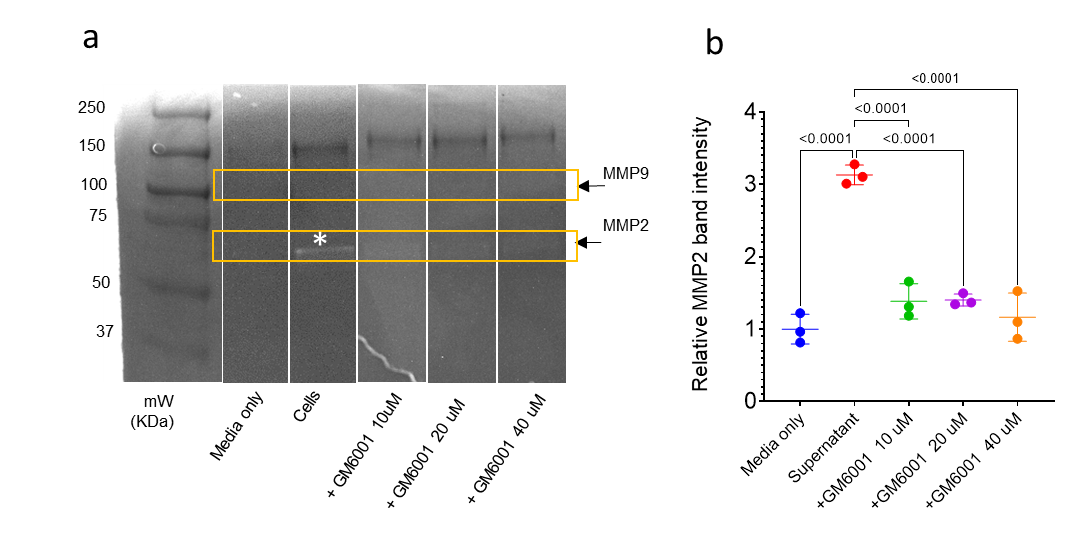

Fig S10: (a) MMPs expression of primary mouse cholangiocytes determined using zymography. Only the MMP2 band was detected (marked with *) suggesting that primary mouse cholangiocytes mostly express MMP2. The MMP2 band was not detected following GM6001 treatment, indicating its effectiveness at inhibiting MMP2-expression. (b) Relative MMP2 band intensity analyses show a significant reduction in MMP2 band intensity after GM6001 treatment. n=3 independent experiments, mean ± S.D. One way ANOVA followed by Dunnett’s post hoc analysis.

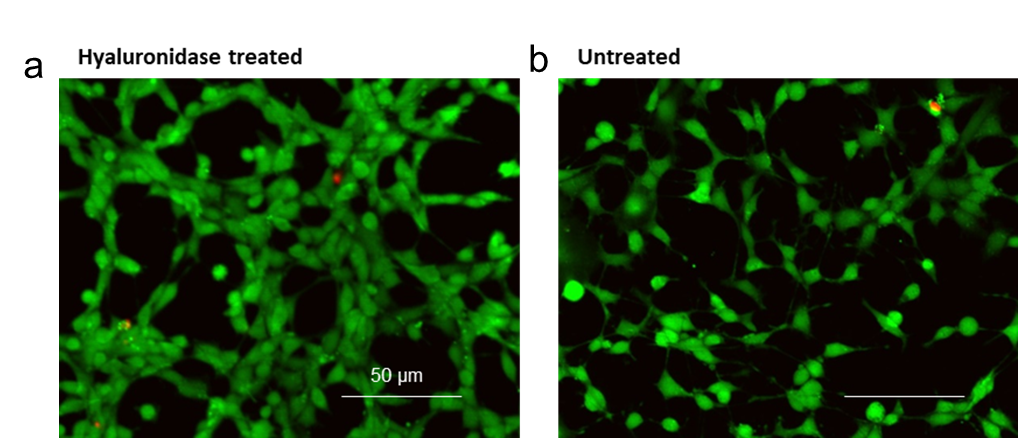

Fig S11: Calcein AM (green, live cells) and Ethidium bromide (red, dead cells) staining of mouse cholangiocytes incubated overnight with either (a) hyaluronidase (2500 U/mL^-1^) or (b) untreated.

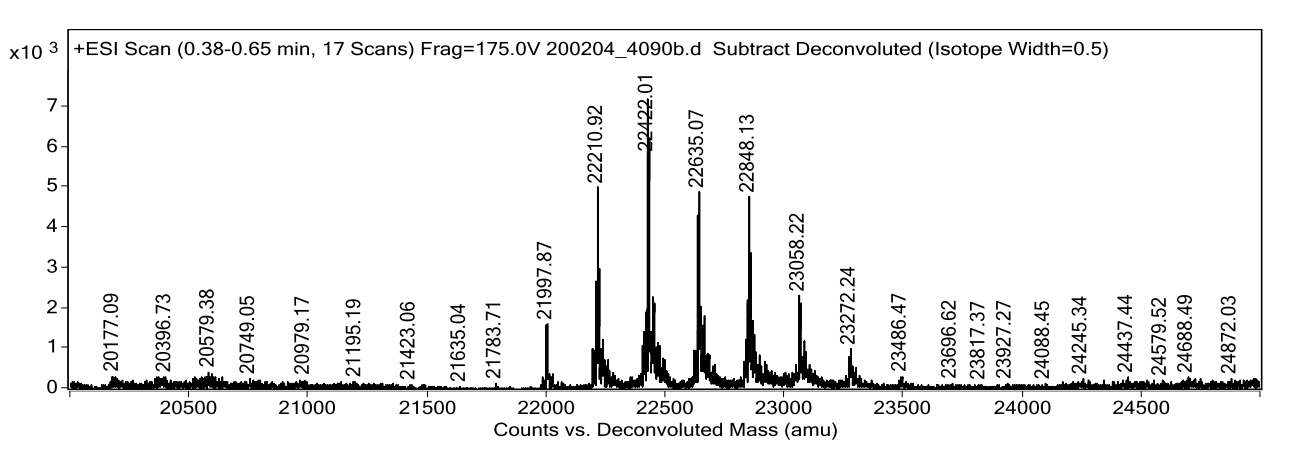

+1 Tz

+2

+3

+4

+5

+6

+7

Fig S12: Mass spectrum of tetrazine modified protein G. Theoretical mass: ~ 21.8 KDa. Observed mass range: 21997.87 to 23272.24 depending on the number of tetrazine groups on Protein G (indicated on the spectrum).

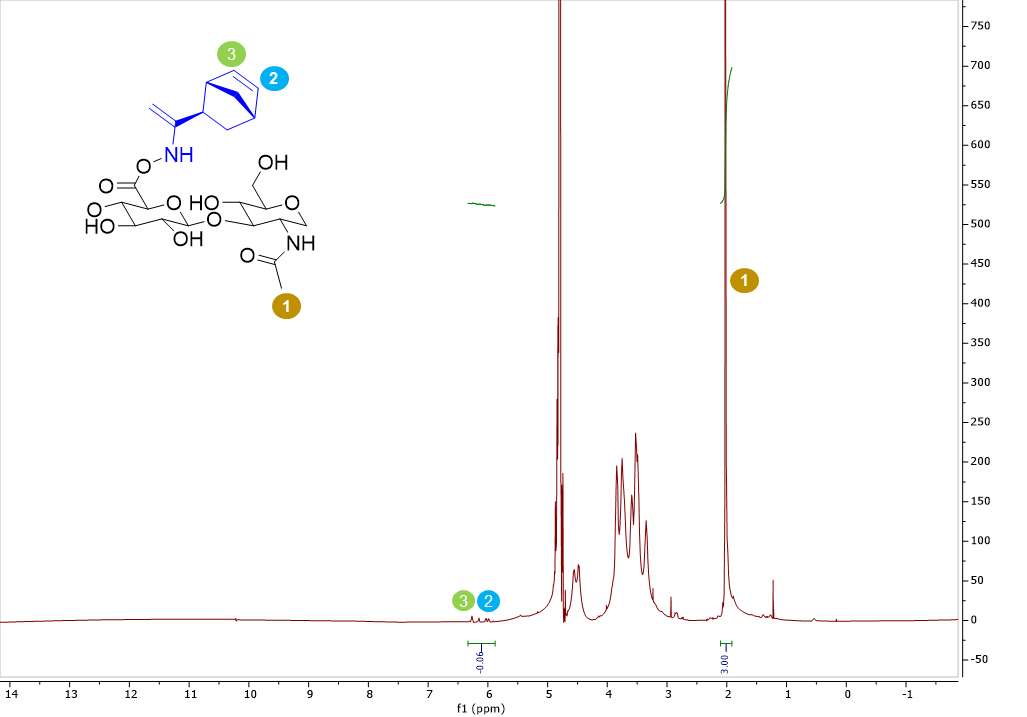

Fig. S13: ^1^H NMR spectrum of norbornene-modified HA in D_2_O. 6% of the carboxlic acids were modified with norborne functional groups. The N-acetyl group of HA (3H at 2 ppm; peak 1) was used as a reference to calculate the degrees of norbornene modification (2H (CH**=**CH) at 6.0–6.4 ppm, Peak 2 and 3). Rest were used for further modification with ketones.

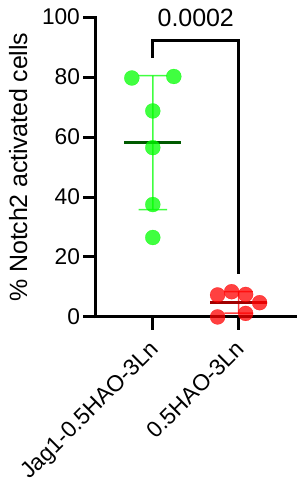

Fig S14: The % CHO reporter cells with Notch2 activation on 0.5HAO-0.3Ln gels with and without immobilized Jag1 in 2D cell culture analyzed at day 2. The % cells with Notch2 activation were determined by counting the cells with fluorescence intensity above fluorescence threshold. The fluorescence threshold was set based on the average fluorescence intensity of CHO cells cultured without Jag1. n=6 independent experiments, mean ± S.D. Student’s t-test.

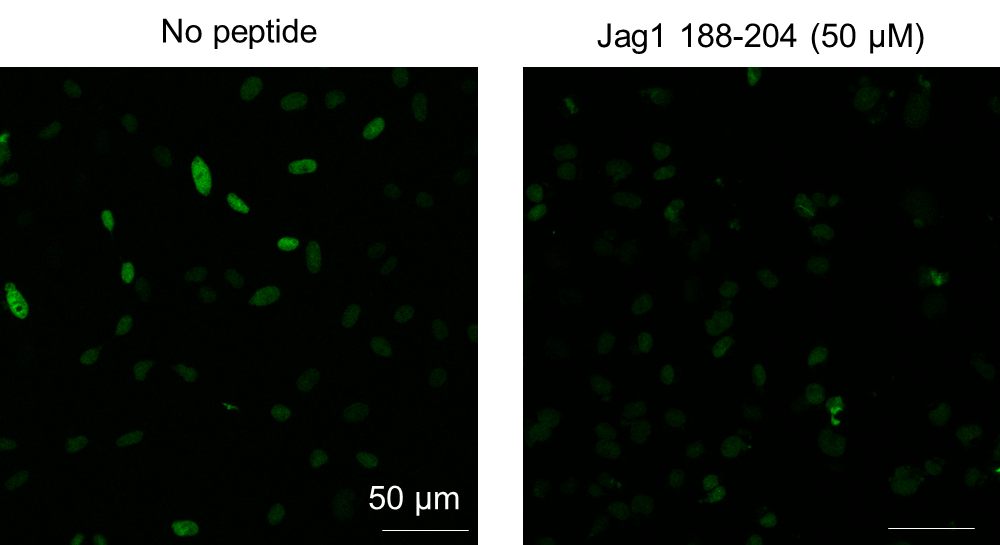

Fig. S15: Jagged1 188-204 peptide does not activate Notch2 signaling using Notch2 fluorescent reporter CHO cells.

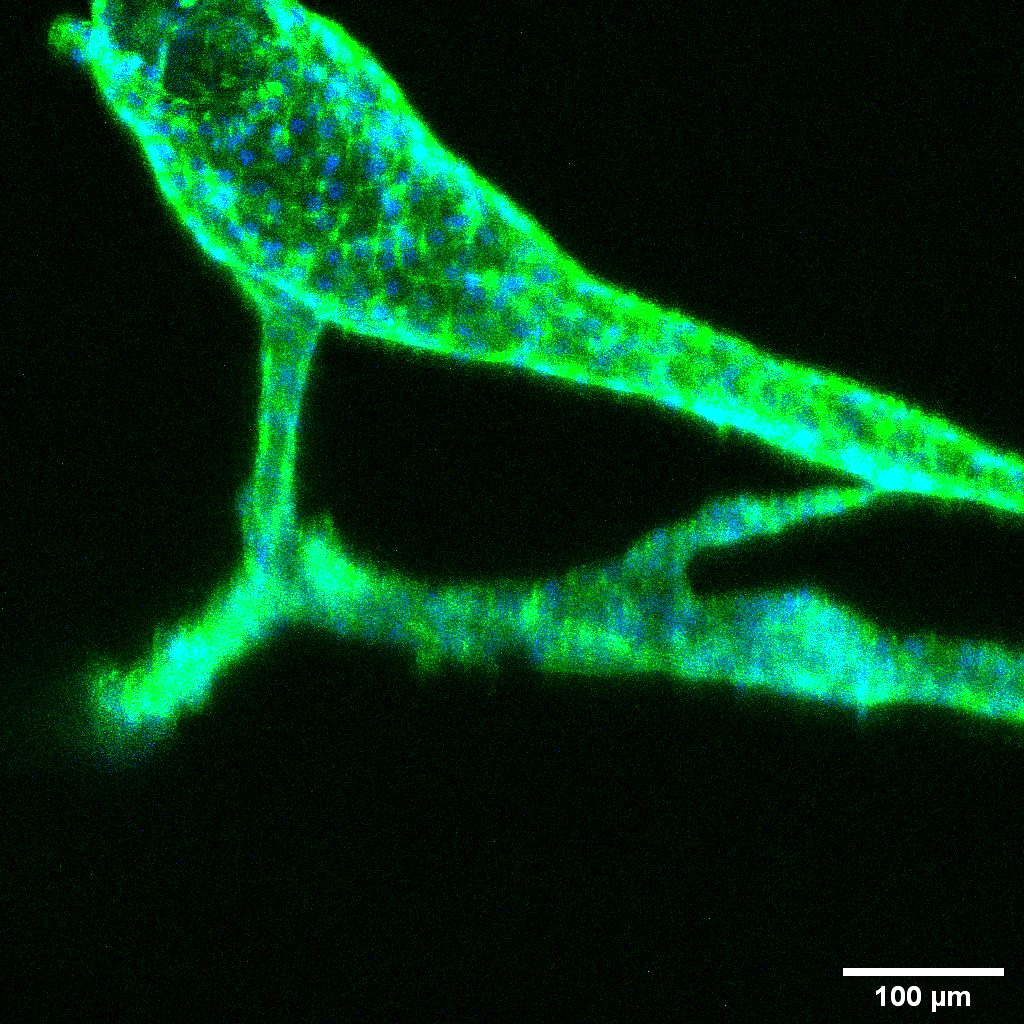

Fig S16. Representative branched ductular structures observed in Jag1-0.5HAO-0.3Ln hydrogels after 14 d of 3D culture. Image represents maximum intensity projection of 36 slices captured with 10 µm step size. Green, F-actin; Blue, Nuclei.

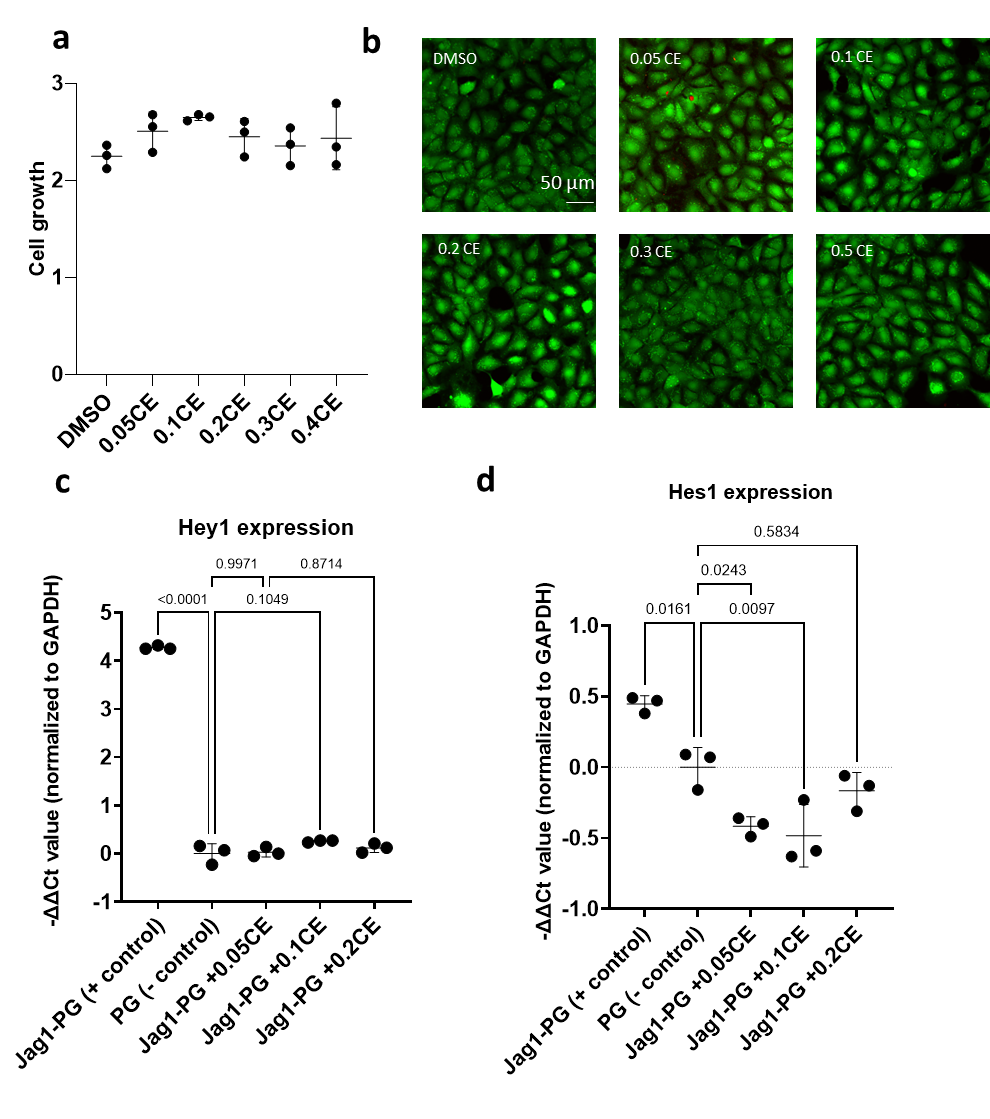

Fig S17: (a) Effect of compound E (Notch inhibitor) on primary mouse cholangiocyte growth measured using PrestoBlue. Data is normalized to day 1. (b) Cholangiocyte viability analyzed using live-dead assay with Calcein AM (green, live cells) and Ethidium bromide (red, dead cells). (c-d) Compound E can successfully inhibit Notch signaling in cholangiocytes as measured by expression analysis of Notch target genes (Hey1, Hes1) after treating cells (cultured on Jag1 immobilized on tissue culture polystyrene) with different concentrations of Compound E for 3 d. The Hey1 and Hes1 expression of treated cells was significantly lower compared to positive control (Jag1 immobilized on polystyrene using PG affinity binding) but statistically indistinguishable from negative control (PG only). PG, protein G.

Table S1: List of primers used for gene expression analysis

| ***SYBR green*** | | |
| --- | --- | --- |
| ***Gene*** | **Forward primer** | **Reverse primer** |
| *Ctgf* | CAAGGACCGCACAGCAGTT | AGAACAGGCGCTCCACTCTG |
| *Cyr61* | AACGAGGACTGCAGCAAAAC | GCGCCATCAATACATGTGCA |
| *Ankrd1* | GGATGTGCCGAGGTTTCTGAA | GTCCGTTTATACTCATCGCAGAC |
| *Areg* | GCTGAGGACAATGCAGGGTAA | GTGACAACTGGGCATCTGGA |
| *Gapdh* | TGTGTCCGTCGTGGATCTGA | TTGCTGTTGAAGTCGCAGGAG |
| **Taqman** | | |
| *Hey1* | Assay ID: Mm00468865_m1 | |
| *Hes1* | Assay ID: Mm01342805_m1 | |
| *Gapdh* | Assay ID: Mm99999915_g1 | |

Table S2: List of primary antibodies and dilutions used

| **Primary antibodies** | **Dilution (in PBS)** | **Vendor and Cat #** |
| --- | --- | --- |
| Rabbit anti cytokeratin 19 (CK19) | 1:100 | Abcam, ab52625 |
| Mouse anti cytokeratin 7 (CK7) | 1:50 | Santa Cruz, sc-23876 |
| Rabbit anti E-Cadherin | 1:200 | Cell signaling Technologies, 3195 |
| Mouse anti β-catenin | 1:50 | Santa Cruz, sc-7963 |
| Goat anti ZO1 | 1:100 | Abcam, ab190085 |
| Rabbit anti HNF4-alpha | 1:2000 | Abcam, ab201460 |
| Mouse anti albumin | 1:50 | Santa Cruz, sc-271605 |
| Secondary antibodies |  |  |
| Goat anti rabbit Alexa Fluor 488 | 1:500 | Thermofisher Scientific, A11008 |
| Goat anti mouse Alexa Fluor 568 | 1:250 | Thermofisher Scientific, A11004 |
| Goat anti mouse Alexa Fluor 488 | 1:500 | Thermofisher Scientific, A11001 |
| Donkey anti goat AlexaFluor 555 | 1:500 | Thermofisher Scientific, A21432 |
